# Pseudoreplication in genomics-scale datasets

**DOI:** 10.1101/2020.11.12.380410

**Authors:** Robin S. Waples, Ryan K. Waples, Eric J. Ward

**Affiliations:** NOAA Fisheries, Northwest Fisheries Science Center 2725 Montlake Blvd. East, Seattle, WA 98112; Department of Biology, Section for Computational and RNA Biology, University of Copenhagen, Copenhagen, Denmark

**Keywords:** degrees of freedom, linkage disequilibrium, *F*_*ST*_, *N*_*e*_, genome size, jackknife variance, simulations

## Abstract

In genomics-scale datasets, loci are closely packed within chromosomes and hence provide correlated information. Averaging across loci as if they were independent creates pseudoreplication, which reduces the effective degrees of freedom (*df’*) compared to the nominal degrees of freedom, *df*. This issue has been known for some time, but consequences have not been systematically quantified across the entire genome. Here we measured pseudoreplication (quantified by the ratio *df’*/*df*) for a common metric of genetic differentiation (*F*_*ST*_) and a common measure of linkage disequilibrium between pairs of loci (*r*^2^). Based on data simulated using models (*SLiM* and *msprime*) that allow efficient forward-in-time and coalescent simulations while precisely controlling population pedigrees, we estimated *df’* and *df’*/*df* by measuring the rate of decline in the variance of mean *F*_*ST*_ and mean *r*^2^ as more loci were used. For both indices, *df’* increases with *N*_*e*_ and genome size, as expected. However, even for large *N*_*e*_ and large genomes, *df’* for mean *r*^2^ plateaus after a few thousand loci, and a variance components analysis indicates that the limiting factor is uncertainty associated with sampling individuals rather than genes. Pseudoreplication is less extreme for *F*_*ST*_, but *df’*/*df* ≤0.01 can occur in datasets using tens of thousands of loci. Commonly-used block-jackknife methods consistently overestimated var(*F*_*ST*_), producing very conservative confidence intervals. Predicting *df’* based on our modeling results as a function of *N*_*e*_, *L*, *S*, and genome size provides a robust way to quantify precision associated with genomics-scale datasets.

## 1 INTRODUCTION

It is now relatively easy to generate data at tens or hundreds of thousands of single-nucleotide polymorphism (SNP) markers for non-model species, even in the absence of a reference genome (da Fonseca et al. 2016; Van Wyngaarden et al. 2017; Aguirre et al. 2019; Choquet et al. 2019; Minias et al. 2019). This opens up vast new opportunities for researchers but also creates a host of analytical challenges, including ascertainment bias (Rosenblum and Novembre 2007; Albrechtsen et al. 2010), phylogenetic inference (Leaché et al. 2015), genotyping errors and missing data (Gautier et al. 2013; Graffelman et al. 2015; Huang and Knowles 2014), and effects of selection (Foll and Gaggiotti 2008; Wolf and Ellegren 2017).

One topic that has not received sufficient attention with respect to genomics data for non-model species is pseudoreplication—lack of independence among datapoints that reduces the total information content. Statistical inference in biology is challenging because biological systems are complex, variable, and subject to measurement and sampling errors. Replication is generally necessary to ensure that apparently-interesting results are not due to small sample sizes and chance. But true replication (which produces multiple independent datapoints) is difficult to achieve, and flaws of experimental design and/or statistical inference that lead to pseudoreplication and overly-optimistic estimates of statistical significance have been found to be widespread in ecology and evolutionary biology (Hurlbert 1984; Ramage et al. 2013; Aarts et al. 2014; Colegrave and Ruxton 2018; Lin et al. 2019).

In the present context, we are interested in use of genetic data to draw inferences about key genetic parameters in real populations of non-model species. For decades, most studies were limited to a few dozen genetic markers, and it was routinely assumed that each marker was independent—or if not, that departures from independence were small enough to be safely ignored. This assumption is no longer tenable in the age of genomics. Most species have at most a few dozen chromosomes (Table 1), so in contemporary datasets a typical chromosome contains thousands of markers—a situation which guarantees that pseudoreplication occurs.

**Table 1.** Summary of information on haploid (1*n*) chromosome number and mean chromosome size and genome size for major taxonomic groups (data from Li et al. 2011).

|  | Chromosome number |  |  |  | Size (Mb) |  |
| --- | --- | --- | --- | --- | --- | --- |
|  | Mean | Min | Max | sd | Chromosome | Genome |
| Prokaryotes | 2.2 | 2 | 4 | 0.50 | 2.5 | 5.7 |
| Unicellular eukaryotes | 16.6 | 2 | 35 | 9.33 | 1.7 | 23.6 |
| Invertebrates | 11.1 | 2 | 16 | 5.18 | 21.7 | 168.4 |
| Vascular plants | 13.4 | 5 | 20 | 5.66 | 47.7 | 558.0 |
| Vertebrates | 25.2 | 8 | 38 | 6.27 | 85.4 | 1933.1 |

Here we focus on two widely-used genetic metrics: *F*_*ST*_, a measure of differentiation among populations; and *r*^2^, a measure of linkage disequilibrium (LD) at pairs of loci. Inter-locus sampling variances for both of these metrics are large, so a high degree of replication is required to obtain reliable estimates of the mean. Let 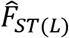 be the estimate of mean *F*_*ST*_ based on data for *L* diallelic loci, and let 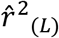 be the two-locus analogue for LD based on *n* = *L*(*L*-1)/2 pairs of *L* loci, where the ‘^’ indicates an estimate. In statistical theory, if one has *k* independent observations of a random variable *x* with standard deviation σ, the standard error of the estimate of mean(*x*) is 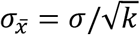. The genetic metrics of interest here also have the property that their variances are proportional to 1/*k*, assuming the data points are independent (Lewontin and Krakauer 1973; Hill 1981).

In population genetic studies, two general sources of replication are available: one can sample multiple individuals from each population, and one can sample multiple loci from each individual. Often the number of individuals that can be sampled is limited by population size or logistical constraints, in which case the only feasible method to increase precision is by increasing the number of loci. Easy access to genomics-scale datasets has made it possible to vastly increase the number of loci sampled per individual—but to what extent do these genes provide independent information about population-level parameters of interest?

We encounter two kinds of pseudoreplication we need to be concerned with. First, if loci do not freely recombine, information they provide is correlated, so (for example) 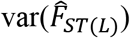 does not decline as fast with addition of more loci as would be the case if the loci were independent. The total information content of a dataset is constrained by the amount of recombination. This constraint means, for example, that even with many millions of SNPs available in human genetic datasets, it is not possible to confidently resolve distant familial relationships with genetic data alone (Thompson 2013). Pseudoreplication due to lack of independent assortment applies both to 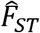 and 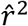. Analyses of LD also generate a second kind of pseudoreplication, caused by overlapping pairs of loci. The approximately *L*^2^/2 different pairs of loci are not independent because each locus occurs in *L*-1 pairwise comparisons.

Problems related to this lack of independence have not been quantified in any systematic way for non-model species. For such species, a typical experimental design involves orders of magnitude more SNP loci than individuals, and if any detailed genomic mapping information is available, loci typically can only be placed on short genomic scaffolds rather than full chromosomes. In this study we investigate how much precision is reduced by pseudoreplication, compared to what it would be if the assumption of independence were completely satisfied. Assuming one has data for *L* diallelic (SNP) loci, the number of datapoints (and the nominal degrees of freedom, *df*, associated with the overall estimate) is *L* for *F*_*ST*_ and *n* ≈ *L*^2^/2 for the LD analyses. Pseudoreplication causes the actual (effective) *df’* (*L*’ or *n*’) to be less than the total number of datapoints (Cox 1984; Giesbrecht 2006). The key question thus can be framed as follows: How much smaller is the effective *df* than the nominal *df*, and how does the *df’/df* ratio depend on aspects of experimental design (number of individuals (*S*) and loci sampled) and uncontrollable parameters of the population(s) of interest (genome size or number of chromosomes, *C*, and effective population size, *N*_*e*_)? The ability to estimate *L*’ or *n*’ would allow an unbiased evaluation of precision and facilitate placing accurate confidence bounds on estimates of key population genetic parameters.

For species (including humans) with reference genomes and linkage map, these dependencies have been addressed to some extent, particularly with respect to multiple testing (Nyholt 2004; Pe’er et al., 2008; Galwey 2009). A popular approach is to use a weighted, block-jackknife that breaks up the genome into blocks of contiguous loci and leaves each block out in jackknife fashion (Busing et al. 1999). LD pruning (Purcell et al. 2007)—excluding loci to reduce LD—can also be used to generate sets of loci that act more independently and thus reduce prereplication. These approaches, however, require detailed mapping information, and even so they only deal with correlations of the original variables. To evaluate the second type of pseudoreplication noted above, it is necessary to consider second-order correlations—that is, the degree to which *r*^2^ (locus 1 × locus 2) is correlated with *r*^2^ (locus 1 × locus 3) and *r*^2^ (locus 2 × locus 3). In theory *n*’ could be calculated this way, but it would require one to specify the relevant covariance matrix, and with ≈*L*^2^/2 pairwise correlations of loci the covariance matrix of these correlations has ≈*L*^4^/8 elements. For a dataset with 1.5 million SNPs, calculating *n*’ therefore would require one to specify over 6.3×10^23^ elements in the covariance matrix—more than Avogadro’s number! Not surprisingly, we are not aware of any attempts to do this.

The objectives of this paper are to develop model-based approximations of *df’*, and to provide general guidance – given known or measurable covariates (*C*, *L*, *S*, *N*_*e*_). Our approach to quantifying the degree of pseudoreplication involves simulating a large number of replicate populations, and for each replicate we calculate mean values for both of the genetic indices (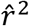, 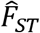). Observed variances of the multilocus metrics across replicates allow us to calculate the *df’* associated with samples of individuals and gene loci. This process was repeated for several evolutionary scenarios involving different combinations of *C* and *N*_*e*_.

We have the following general expectations:

1. As more loci are packed into a fixed number of chromosomes, the number of loci per chromosome increases, minimum distance between a new locus and existing loci shrinks (Figure S1), and the marginal increase in information content provided by each new locus declines. Therefore, we expect that as the ratio *L*/*C* increases, pseudoreplication will increase and the ratio of effective to nominal df will decrease for both metrics. In addition, the rate of decay of LD with distance between loci increases with effective population size (Figure S2). So, we expect that, for a given *L*/*C* ratio, *n*’ will be smaller for populations with smaller *N*_*e*_. These effects, however, have not been quantified across the entire genome.
2. It seems intuitive that the magnitude of LD increases with the number of loci, but that is not actually the case. Each new locus increases opportunities for that locus to be in LD with existing loci, but this is balanced by new pairwise comparisons with loci on different chromosomes. As a result, the probability that any two randomly-chosen loci will be in any particular LD association is independent of the number of loci, but it does depend on genome size and *N*_*e*_ (Waples et al. 2016). Therefore, we expect that the first type of pseudoreplication for LD analyses will be inversely correlated with both *N*_*e*_ and *C*. The second type of pseudoreplication arising from LD analyses—that caused by overlapping pairs of the same loci—has received little study. However, we note that, of the ~*L*^2^/2 pairwise comparisons of *L* loci, only *L*/2 are completely independent (non-overlapping), so the proportion of independent comparisons is 1/*L*. Hence, we expect that this type of pseudoreplication will increase with the number of loci.

We hope to find one or both of the following:

a. Patterns in empirical variances across simulated datasets that allow us to provide general guidance for users interested in predicting *df’*, based on measurable or estimable covariates (*C*, *L*, *S*, *N*_*e*_).
b. Existing jackknife methods prove to be reliable at estimating precision for a given dataset.

## 2 METHODS

Below we provide an overview of methods used; for more details, see Supporting Information. Table 2 summarizes notation.

**Table 2.**
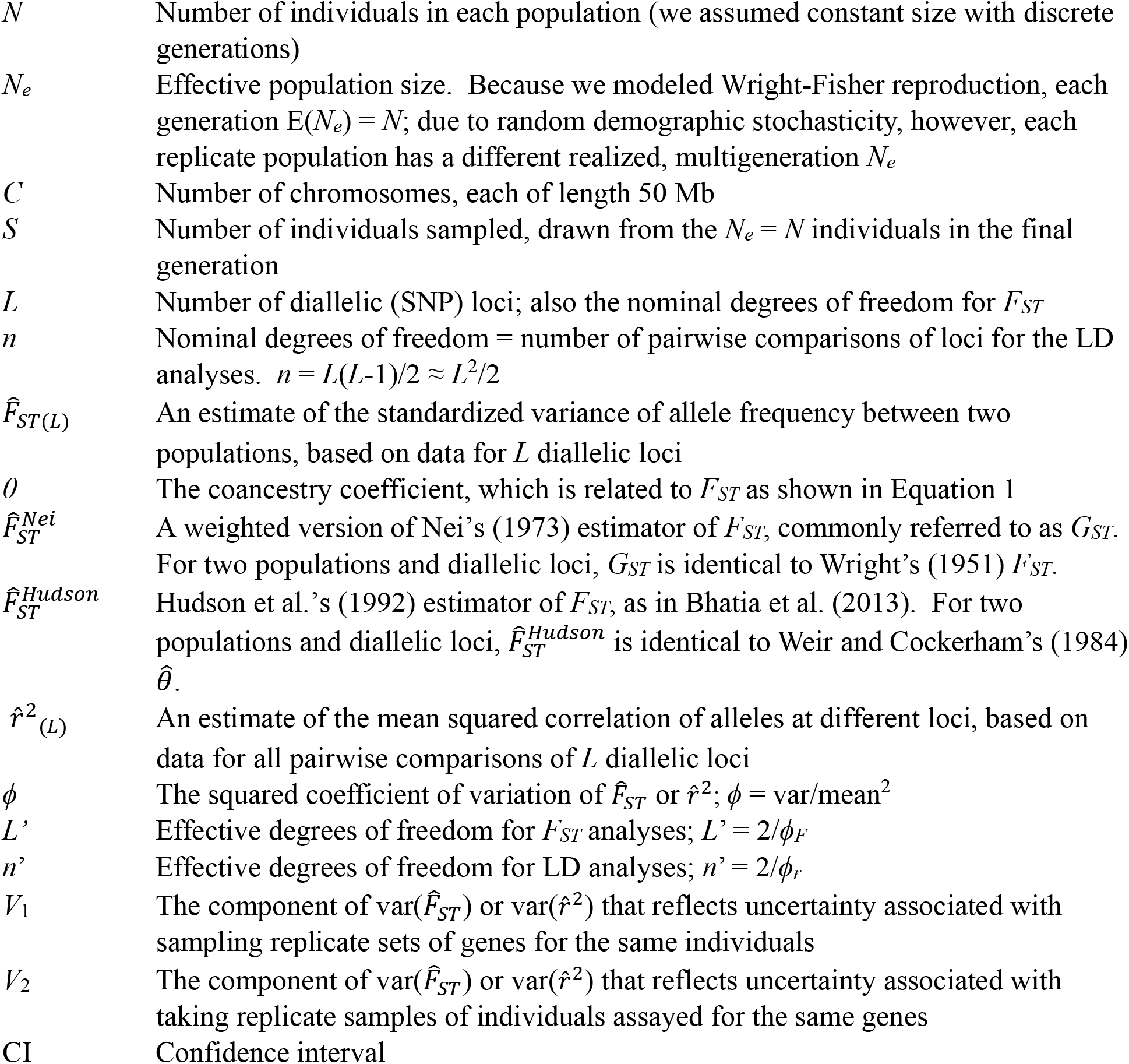
Notation used in this study

### 2.1 Conceptual Framework

Our focus is on actual populations, each of which has a single, realized population pedigree (Wakeley et al. 2012; Ralph 2019). To mimic sampling of individuals and genes from real populations, we combine efficient coalescent and forward simulation programs (*SLiM*, Messer 2013; and *msprime*, Kelleher et al. 2016) that allow control over multigenerational pedigrees in replicate populations (Haller et al. 2019). Our experimental design explicitly models two major sources of uncertainty in estimating population-level parameters: sampling of individuals and sampling of genes. Both processes are important when evaluating uncertainty around parameter estimates. If replication only occurs across genes, estimates will converge on values determined by the pedigree of the observed individuals (Waples and Faulkner 2009; Wakeley et al. 2012; King et al. 2018). In general, however, one wants to draw inference about the entire population, and uncertainty related to sampling individuals from the population cannot be eliminated by intensive replication across genes.

### 2.2 Modeled Scenarios and Simulations

For 16 different evolutionary scenarios (combinations of effective size [*N*_*e*_=50,200,800,3200] and number of chromosomes [*C*=1,4,16,64]), we simulated four separate ancestral populations of *N=N*_*e*_ diploid individuals for 10*N*_*e*_ generations under Wright-Fisher (WF) reproduction and ensured that each ancestral population fully coalesced. The simulations modeled the process of recombination and mutation in realistic-sized genomes with distinct chromosomes. Genomes were simulated with varying numbers of chromosomes [1, 4, 16, 64], each of which was 50 Mb (5×10^7^ bp) long, with a recombination rate of 10^−8^ per bp per generation. Each ancestral population then split into four daughter populations of size *N*_*e*_, which evolved in isolation for enough generations that expected *F*_*ST*_ ≈0.05-0.1, common values for natural populations of many species. From this point, the models differed slightly for analyses of *F*_*ST*_ and LD.

In the LD analyses, for each of the 4×4=16 population pedigrees, we created eight non-overlapping sets of loci by adding mutations to gene trees present in the population pedigree (Figure 1). Gene trees reflect the evolutionary history of genes and haplotypes back in time. For each scenario, the 4×4×8 hierarchical design produced 128 replicate populations with up to 75K diallelic loci (up to 100K loci for *N*_*e*_=3200). From each replicate population, we took four random subsamples of *S*=25,50,100 diploid individuals (only *S*≤50 for *N*_*e*_=50), and data for each subsample were analyzed for *L*=100-75,000 loci. Except for *N*_*e*_=3200, we also took exhaustive samples of the population (*S*=*N*_*e*_). This experimental design allowed us to calculate two separate variance components: *V*_1_ (same individuals, different loci) and *V*_2_ (same loci, different but potentially overlapping samples of individuals) (Table 3). Data from cells on the diagonal (different individuals and different loci) were used to calculate *df’* for the LD analyses (*n*’), as described below.

**Figure 1.**
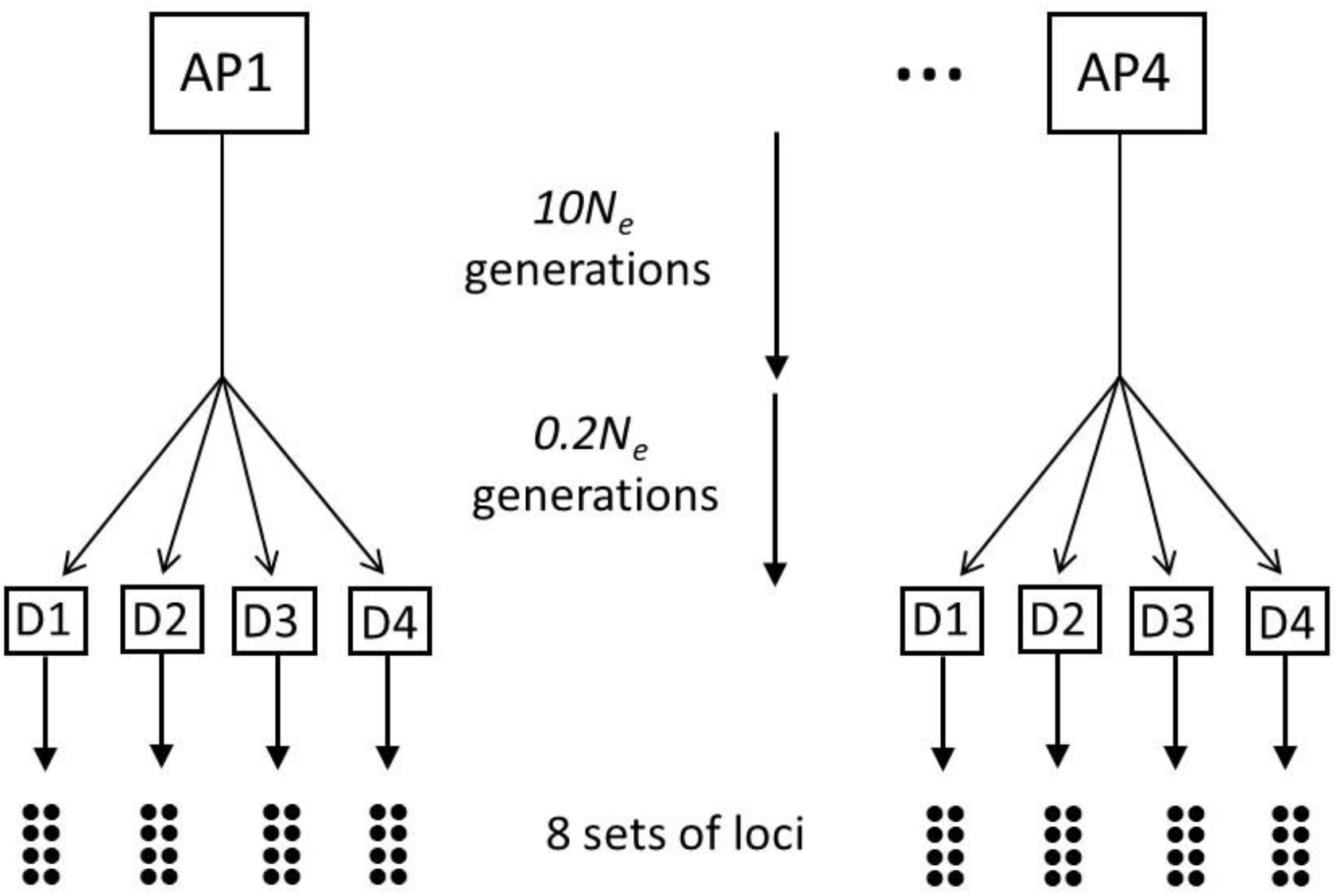
Experimental design for simulations. For each evolutionary scenario (combination of *N*_*e*_ and genome size), four ancestral populations (AP1–AP4) were simulated to ensure coalescence (10*N*_*e*_ generations), at which point each ancestral population split into four daughter populations (D1-D4). The 4×4 = 16 daughter populations then evolved independently under isolation for *t* = 0.2*N*_*e*_ generations. Subsequently, the model differed slightly for the *F*_*ST*_ and LD analyses. In the latter (as depicted in the figure), for each daughter population, eight mutational replicates (different set of loci) were generated based on the same pedigree, producing a total of 128 replicates for each evolutionary scenario. For *F*_*ST*_, each set of four daughter populations allowed six pairwise comparisons of populations, and for each two-population pedigree six mutational replicates were generated (Figure S4).

**Table 3.**
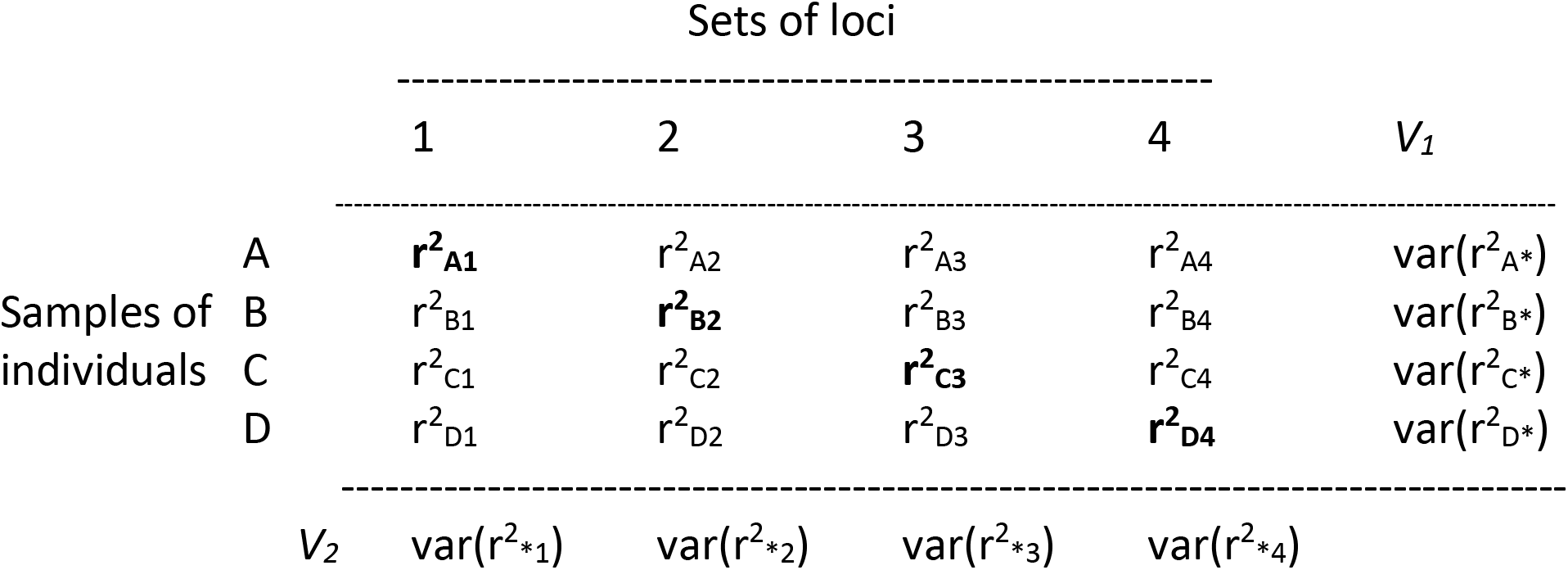
Experimental design for sampling individuals and genes for analyses of LD. From each population, potentially-overlapping subsets of individuals (rows) were drawn from the *N*_*e*_ total individuals in the final generation. Subsets of *L* loci (columns) were non-overlapping. Mean *r*^*2*^ was calculated for each cell. The variance among mean *r*^*2*^ within rows was used to estimate variance component *V*_*1*_ (same individuals, different loci), and the variance within columns was used to estimate variance component *V*_*2*_ (same loci, different individuals). A measure of pseduoreplication, *ϕ*_*r*_, was calculated across mean *r*^*2*^ for the cells (in **bold**) along the diagonal (different sets of individuals and loci). This sampling design was repeated for different numbers of loci (*L*), sampled individuals (*S*), chromosome number (*C*), and *N*_*e*_. See the text and Figure S3 for details of a modified sampling design for *F*_*ST*_ that involved pairwise comparisons of daughter populations.

In the *F*_*ST*_ analyses, for each of the four ancestral populations, the four daughter populations shown in Figure1 allowed 6 pairwise comparisons among populations, for a total of 24 two-population pedigrees (Figure S3). Six mutational replicates of 200K loci were created for each pedigree, and eight samples for each of several different sizes were taken from each mutational replicate.

The resulting distributions of minor allele frequency (MAF) followed the familiar U-shaped pattern expected at mutation-drift equilibrium (Figure S4). In addition to the *SLiM*/*msprime* simulations that organize the genome into chromosomes, for LD we also modeled scenarios with unlinked loci and infinite *N*_*e*_ to provide insights into asymptotic behavior.

### 2.3 Genetic Indices

For each replicate in each evolutionary scenario, we calculated 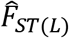 and 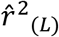 across variable numbers of loci or locus pairs. LD analyses were conducted using both all pairs of loci, and only pairs on different chromosomes. For each pair, the sample estimate of the squared correlation coefficient 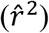 was computed using the Pearson product-moment correlation of diploid genotypes. After applying the sample-size adjustments implemented in *LDN*_*E*_ (Waples and Do 2008), mean 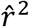 can be used to estimate *N*_*e*_. Because this method assumes all loci are unlinked, use of all locus pairs leads to a predictable pattern of downward bias in 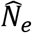 that is inversely proportional to log(*C*) (Waples et al. 2016), whereas 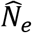 is unbiased when computed only using pairs of loci on different chromosomes (Figure S5). To reduce effects of rare alleles, within each sample we omitted loci with two or fewer copies of the minor allele.

We used two methods for estimating the standardized variance of allele frequency among populations, *F*_*ST*_. Nei’s (1973) gene diversity method calculates *G*_*ST*_ = (*H*_*T*_-*H*_*s*_)/*H*_*T*_, where *H*_*s*_ is the expected within-population heterozygosity (averaged across both populations) and *H*_*T*_ is expected total heterozygosity, based on mean allele frequencies across both populations, without an adjustment for sample size. Calculated this way, and with just two alleles at each locus, *G*_*ST*_ is identical to Wright’s (1951) *F*_*ST*_ (Nei and Chakravarti 1977); we refer to this estimator as 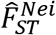.

Another widely-used measure of population differentiation is the coancestry coefficient, *θ* (Cockerham 1969; Reynolds et al. 1983); the relationship between the two parameters is given by (Cockerham and Weir 1987)

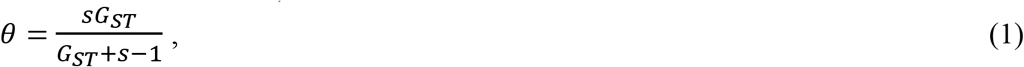

where *s* is the number of subpopulations. For *s*=2 (as considered here) this reduces to *θ=*2*G*_*ST*_/(*G*_*ST*_+1), which is close to 2*G*_*ST*_ if *G*_*ST*_ is small. We considered Hudson et al.’s (1992) estimator 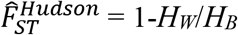, where *H*_*W*_ and *H*_*B*_ are the mean numbers of allelic differences within and between populations, respectively; the formula we used was Equation 10 in Bhatia et al. (2013). Under conditions modeled here (two populations, equal sample sizes), 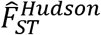 is identical to Weir and Cockerham’s (1984) 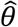, or nearly so (Bhatia et al. 2013).

Because performance of multilocus *F* statistics is generally better when they are computed as ratios of multi-locus means rather than means of single-locus ratios (Jorde and Ryman 2007; Bhatia et al. 2013), we evaluated both methods for calculating 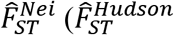 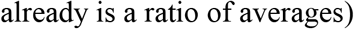. We considered three ascertainment schemes for identifying variable loci: 1) loci variable in at least one of the two populations; 2) loci variable in the ancestral population; 3) loci with overall MAF≥0.05 in the two samples combined.

### 2.4 Effective Degrees of Freedom and Effective Number of Loci

For analysis of population differentiation, effective *df* was calculated from observed variances of 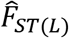 using the relationship 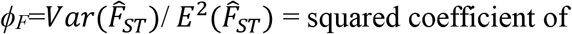 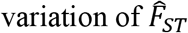 (Lewontin and Krakauer 1973). If 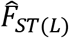 is based on *L* independent diallelic loci, the following relationship should hold: *ϕ*_*F*_=2/*L*. Therefore, we computed *df’* from the empirical variance of 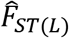 as *L’*=2*ϕ*_*F*_. *L’* can be interpreted as the number of independent loci that would be expected to produce the value of *ϕ*_*F*_ observed in the data (Cox 1984; Giesbrecht 2006). The ratio *L*’/*L* therefore provides an index of how much nonindependence has increased variance of the estimator.

Many commonly-used estimators of *F*_*ST*_ and related quantities (Weir and Cockerham 1984; Hudson et al. 1992; Patterson et al. 2012) include adjustments for sampling individuals and/or populations, and this complicates comparisons with theoretical expectations of the variance-to-mean ratio that depend on raw *F*_*ST*_ values. This is why, for estimating *L*’, we used Nei’s (1973) gene diversity method, which does not include a sample-size adjustment. For Hudson’s estimator, we measured the rate of decline in the variance of 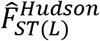 as more loci were used in the analysis, and this provided an alternative way to quantify the degree of pseudoreplication.

Hill (1981) showed that the relationship *ϕ*_*r*_ =2/*n* also applies to 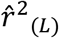, under the assumption that all *n* pairwise comparisons of *L* loci are independent. Accordingly, we estimated *df’* for LD analyses as *n*’ = 2/*ϕ*_*r*_. Because the effective number of pairs of loci is not a very intuitive metric, in some cases (Figures 2 and 3) we presented LD results in terms of the effective number of loci (*L*’), which is the number of loci that would produce the actual observed variance of 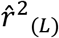, if all the resulting locus pairs were independent. To a good approximation, if *n*’ is the effective number of locus pairs, then the effective number of loci that would produce *n*’ is *L*’ ≈ √2*n*’ [the exact value is 0.5+√(0.25+2*n*’), but this simple approximation is much more intuitive].

**Figure 2.**
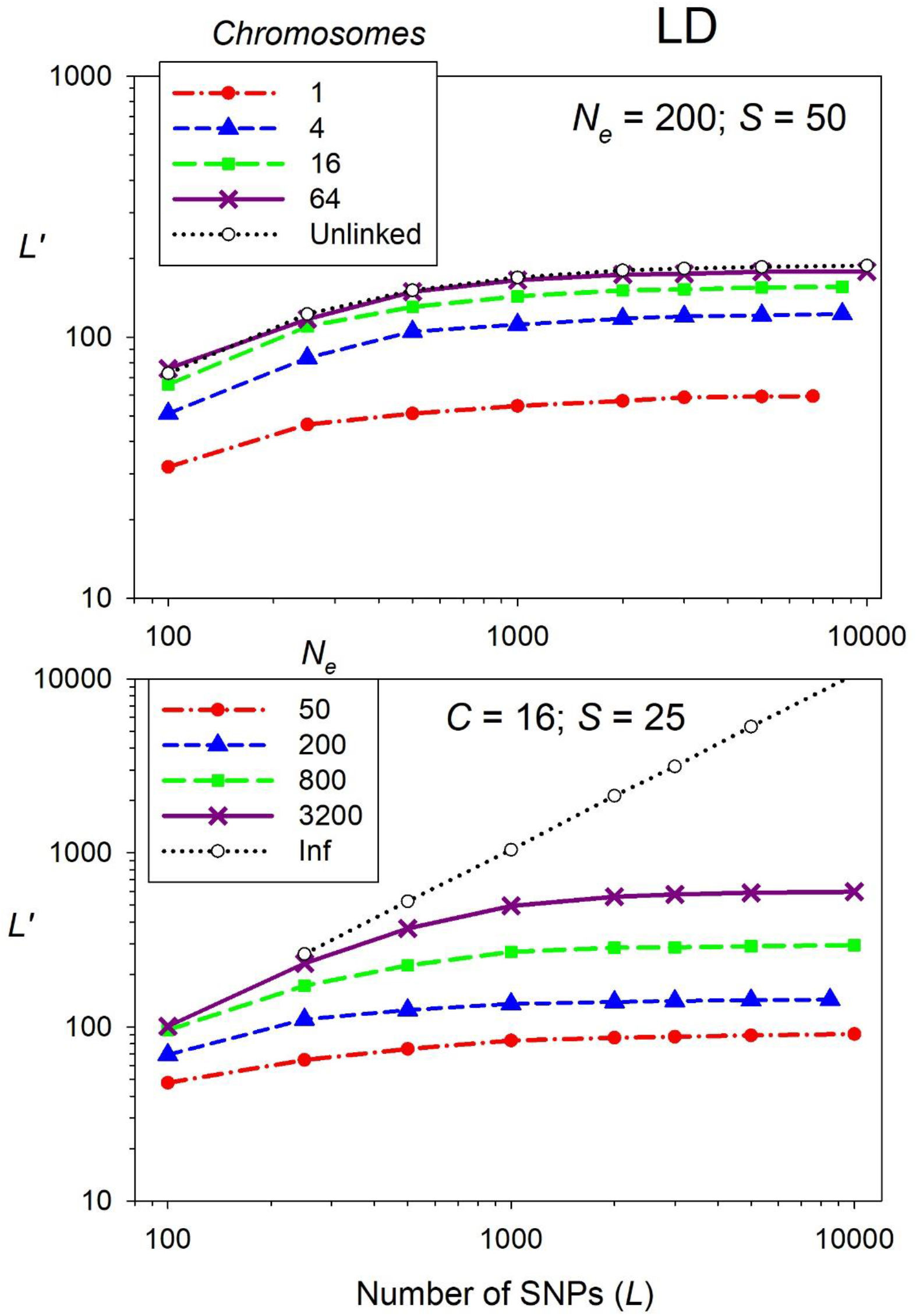
Effective number of loci (*L*’) for mean *r*^2^ as a function of the number of diallelic (SNP) loci, *L*. Top: Influence of number of chromosomes (*C*), with *N*_*e*_ = 200 and *S* = 50. Bottom: Influence of *N*_*e*_, with *C* = 16 and *S* = 25. Mean *r*^2^ was calculated across all *n* = *L*(*L*-1)/2 pairs of loci. Figure S8 (Supplementary Information) shows these same results except the *Y* axis is plotted as the effective number of locus pairs (*n*’).

**Figure 3.**
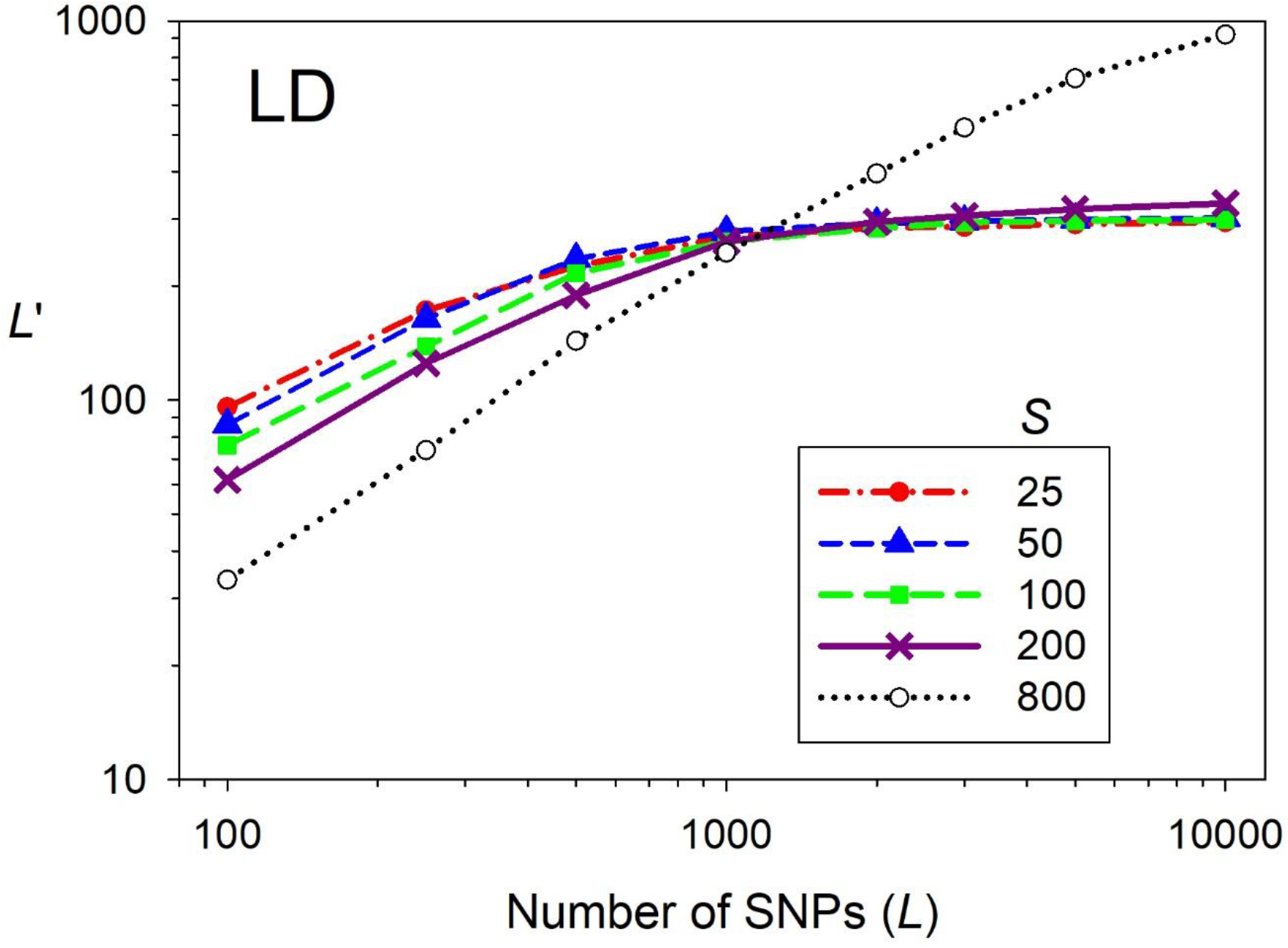
Effective number of loci (*L*’) for mean *r*^2^ as a function of the sample size of individuals (*S* = 25–800) and the number of diallelic (SNP) loci, *L*. Results are for *N*_*e*_ = 800, *C* = 16, and using all pairwise comparisons of loci.

### 2.5 Model Fitting

We considered a wide range of covariates [*N*_*e*_, *C*, *S*, *L*, and (for LD) *n*] as predictors of *df’*. To account for potential non-linearities, we also considered transformed responses of these original variables (Table S1). Because of asymptotic relationships between *df’* and the predictor variables (Figure S6), we focused inference on fitting statistical models with asymptotes, rather than using linear models. The general function we used to parameterize the models was

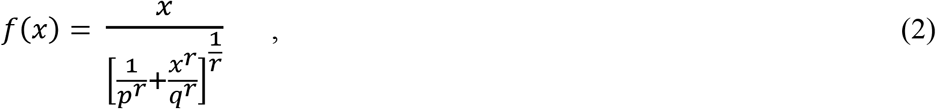

where parameters *p*, *q*, and *r* control the shape of the function. When shape parameter *r*=1, this function takes the form of the familiar Michaelis–Menten equation (also known in fisheries as the Beverton–Holt stock recruitment model; Beverton and Holt 1957). We considered models with both covariates and transformed functions of covariates; these latter models treated each parameter (*p*, *q*, *r*) as linear functions of other predictors; e.g., *p* = *b***X**, where **X** is a design matrix of predictors and *b* are estimated coefficients. All models were fit in R (R Core Team 2020) using maximum likelihood. We used Akaike’s Information Criterion (AIC) to evaluate which combinations of parameters were best supported. Additional details about model fitting are in Supporting Information.

### 2.6 Confidence Intervals

To evaluate accuracy of our estimates of *df’*, for selected scenarios we generated many (>1000) new samples of individuals and loci, and for each sample we calculated 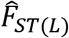 or 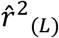 averaged over data for 500–5000 SNPs (for LD) or 5000–50000 SNPs (for *F*_*ST*_). We calculated confidence intervals (CIs) around these means using standard statistical theory (Equations S6 and S7). Width of the CIs was calculated two ways, using: 1) our modeled estimates of *df’*; 2) a published jackknife method (Busing et al. 1999 for *F*_*ST*_, Jones et al. 2016 for 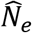 based on *r*^2^).

## 3 RESULTS

### 3.1 Linkage Disequilibrium

The four different ancestral populations produced modest differences in mean *r*^2^ when averaged across all descendant populations, and within each daughter population the eight mutational replicates also produced relatively small differences in mean *r*^2^ (Figure S7). In contrast, daughter populations descended from the same ancestral population varied substantially in the mean magnitude of LD. This result emphasizes the importance of accounting for recent population pedigrees in assessing variability of genetic indices and shows the sensitivity of LD-based estimates of *N*_*e*_ to recent effective population size (Waples and Faulkner 2009).

#### 3.1.1 Effective degrees of freedom

Table S2 gives our estimates of both *n*’ and *L*’ for *r*^2^ for every scenario we simulated. We expected substantial effects of genome size and *N*_*e*_ on *n*’ and *L*’, and those expectations were borne out (Figure 2 shows results for *L*’; see Figure S8 for the same results in terms of *n*’). For a given number of loci, *L*’ increased smoothly as the number of chromosomes increased from 1 to 64, with results for *C* = 64 being largely indistinguishable from those for unlinked loci (Figure 2, top). The influence of effective size was even stronger: *L*’ increased systematically as *N*_*e*_ increased from 50 to 3200 (Figure 2, bottom); however, for large numbers of loci, *L*’ for *N*_*e*_=3200 was still substantially lower than the *L*’≈ *L* that was found for *N*_*e*_=∞. With 5000 loci, var(*r*^2^) decreased by a factor of 5 when *N*_*e*_ increased four-fold from 50 to 200, and the variance decreased by a factor of 68 when *N*_*e*_ increased 64-fold to 3200 (using data from Table S2 with *S*=25 and *C*=4). In contrast, quadrupling the number of chromosomes (from 1 to 4, with *N*_*e*_ fixed at 50) decreased var(*r*^2^) only by 2.5×, and a 64-fold increase in *C* reduced var(*r*^2^) only by 10×.

Relatively speaking, sample size of individuals has less influence on *L*’: with *C*=16, a four-fold increase in *S* reduced var(*r*^2^) by only 1.9× for *N*_*e*_=200 and by only 6% for *N*_*e*_=800. There was one notable exception, however: we found a qualitative difference in *L*’ between scenarios in which the entire population was sampled and those in which only a subset of individuals was analyzed (Figure 3). With exhaustive sampling (*S*=*N*=*N*_*e*_), *L*’ continues to rise, albeit increasingly slowly, as larger and larger numbers of loci are used to compute 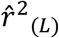. In contrast, for incomplete sampling (*S*<*N*), *L*’ rapidly plateaus once about 1000 loci are used, after which additional loci do little to increase precision. As a consequence, for 10,000 SNPs *L*’ is only about 3% of the number of loci used (and about 3 orders of magnitude smaller than the number of pairwise comparisons; Table S2). Furthermore, as long as the entire population is not exhaustively sampled, sample size has relatively little effect on *L*’ (Figure 3). Samples of individuals were all drawn from the *N*=*N*_*e*_ individuals in the final simulated generation, and the variance associated with replicate hypergeometric samples is proportional to (*N*-*S*)/*N*. *A priori*, therefore, we expected that *L*’ would increase smoothly with sample size. Two factors likely explain the patterns actually observed.

First, larger samples produce less sampling error (Hill 1981; Waples 2006), which reduces both 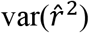 and 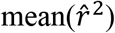. Because *n*’ depends on *ϕ*_*r*_ which is the ratio of 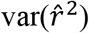 to 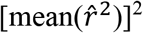, the net effect of increasing *S* is only a modest change in *n*’ and *L*’. Second, the variance components analysis (Figure 4) shows that whereas *V*_1_(variance among replicate sets of loci assayed on the same individuals) continues to decline rapidly with increasing numbers of loci, *V*_2_(variance among different samples of individuals assayed for the same loci) does not decline much after about 500-1000 loci are used. As a consequence, except for small numbers of loci, *V*_2_ dominates overall 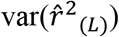 and hence this component largely determines *n*’. Although sampling a larger fraction of the population does reduce *V*_2_, the resulting variance still is much larger than *V*_1_ and still dominates *n*’ and *L*’. Note also that the actual variance of 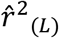 is smaller than the sum of *V*_1_ and *V*_2_ (Figure 4), which indicates that the two variance components must be negatively correlated to some extent. In what follows, we focus on *S*<*N*, which is the most common scenario in studies of natural populations.

**Figure 4.**
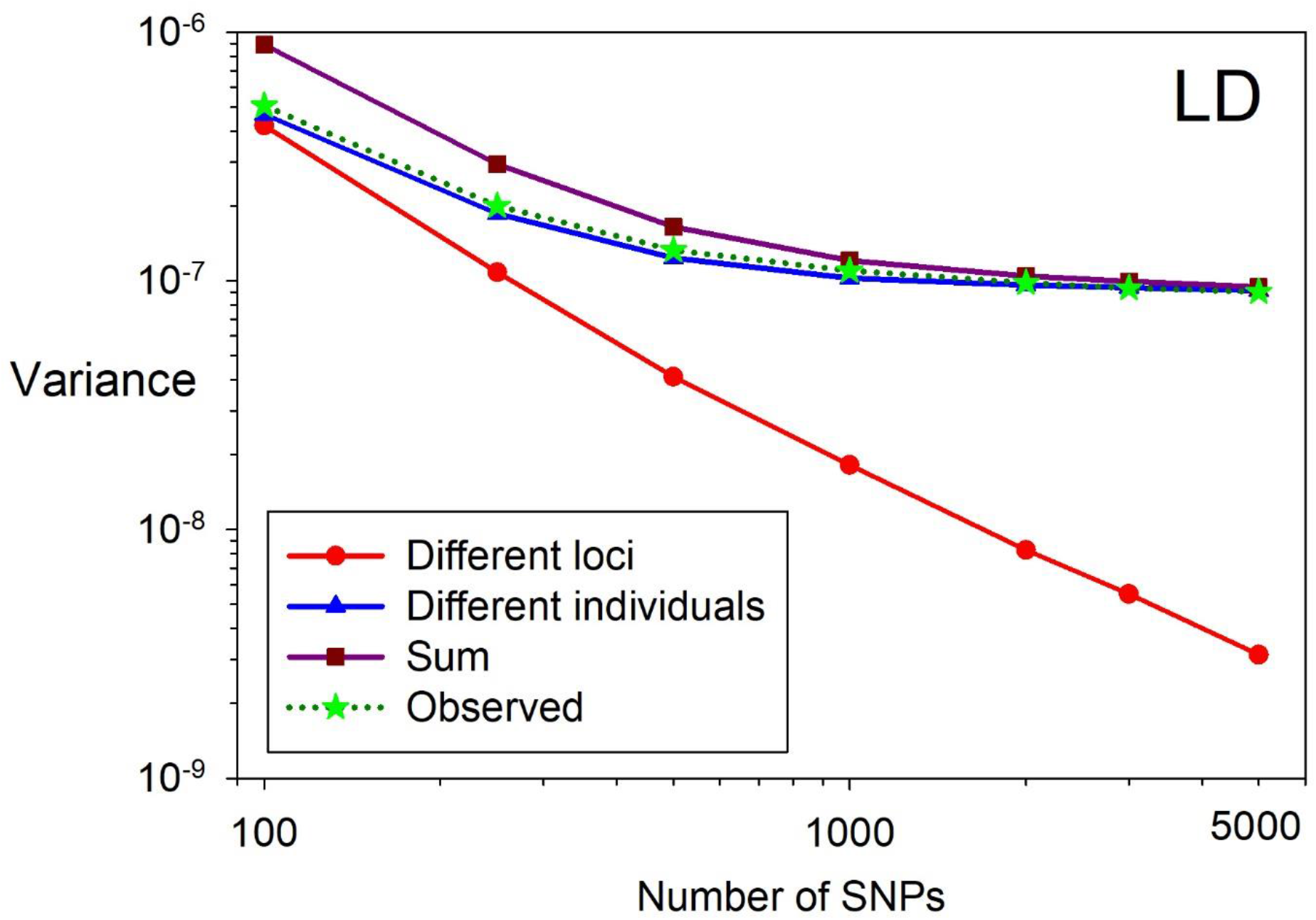
Variance components analysis for mean *r*^2^. As depicted in Table 3, *V*_*1*_ is the variance of mean *r*^2^ for the same individuals assayed for different, non-overlapping sets of loci, and *V*_*2*_ is the variance of mean *r*^2^ for different (potentially overlapping) sets of individuals assayed for the same loci. “Sum” = *V*_*1*_+*V*_*2*_ and “Observed” is the total observed variance of mean *r*^2^. Results are for *N*_*e*_ = 200, *C* = 16, *S* = 50, and using all pairwise comparisons of loci.

Restricting LD analyses to pairs of loci on different chromosomes reduces the number of pairwise comparisons by the proportion 1/*C*, but this has little effect on *n*’, which is essentially the same regardless whether same-chromosome comparisons are allowed or not (Figure S9; the difference is a bit larger for small genomes, where linkage has a stronger effect). Unless otherwise noted, all results presented are for all pairwise comparisons.

The fact that LD results for unlinked loci approximate those for simulations with 64 chromosomes, and that both scenarios show that *L*’ and *n*’ both fail to increase much after a few thousand loci are used (Figures 2 and S8, top), indicate that physical linkage is not the primary factor that reduces *df’* for analyses involving LD. The second factor relevant to two-locus analyses involves overlapping pairs of the same loci. We expect that, if we have genotypes for four unlinked loci (*w,x,y,z*) and compute all six pairwise correlations, the correlation [*w,x*] will be independent of the correlation [*y,z*], but to what extent does the correlation [*w,x*] provide independent information to the correlation [*w,y*], since one locus is shared?

To evaluate this factor, we simulated unlinked loci in many replicate populations and calculated *r*^2^ for each pair of loci in each replicate, across all *N*_*e*_ individuals in the population (see Detailed Methods in Supporting Information). Then, across all replicates, we computed the squared correlation between *r*^2^ values that did and did not share one locus. Results show that correlations between pairs of *r*^2^ values that did not share a locus were essentially 0, regardless how large or small *N*_*e*_ was (Figure S10). This is also the result one obtains if one compares correlations between pairs of vectors of *i.i.d.* random variables, regardless whether the pairwise correlations being compared share one variable or not (data not shown). However, when the data being compared are generated by a process mediated by a pedigree (as occurs during reproduction in finite populations), then the pairwise correlations are not independent when they share one variable, and the degree of non-independence is inversely related to *N*_*e*_ (Figure S10).

#### 3.1.2 Confidence intervals

For datasets with more than 1000 loci, the number of pairwise comparisons of loci is of order 10^6^ or higher, in which case parametric CIs for 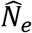 that assume all datapoints are independent become vanishingly small (Figure 5). Based on estimates of *df’* obtained in this study, actual CIs for large numbers of loci are much wider, and this difference is important to understand to avoid misleading conclusions about precision.

**Figure 5.**
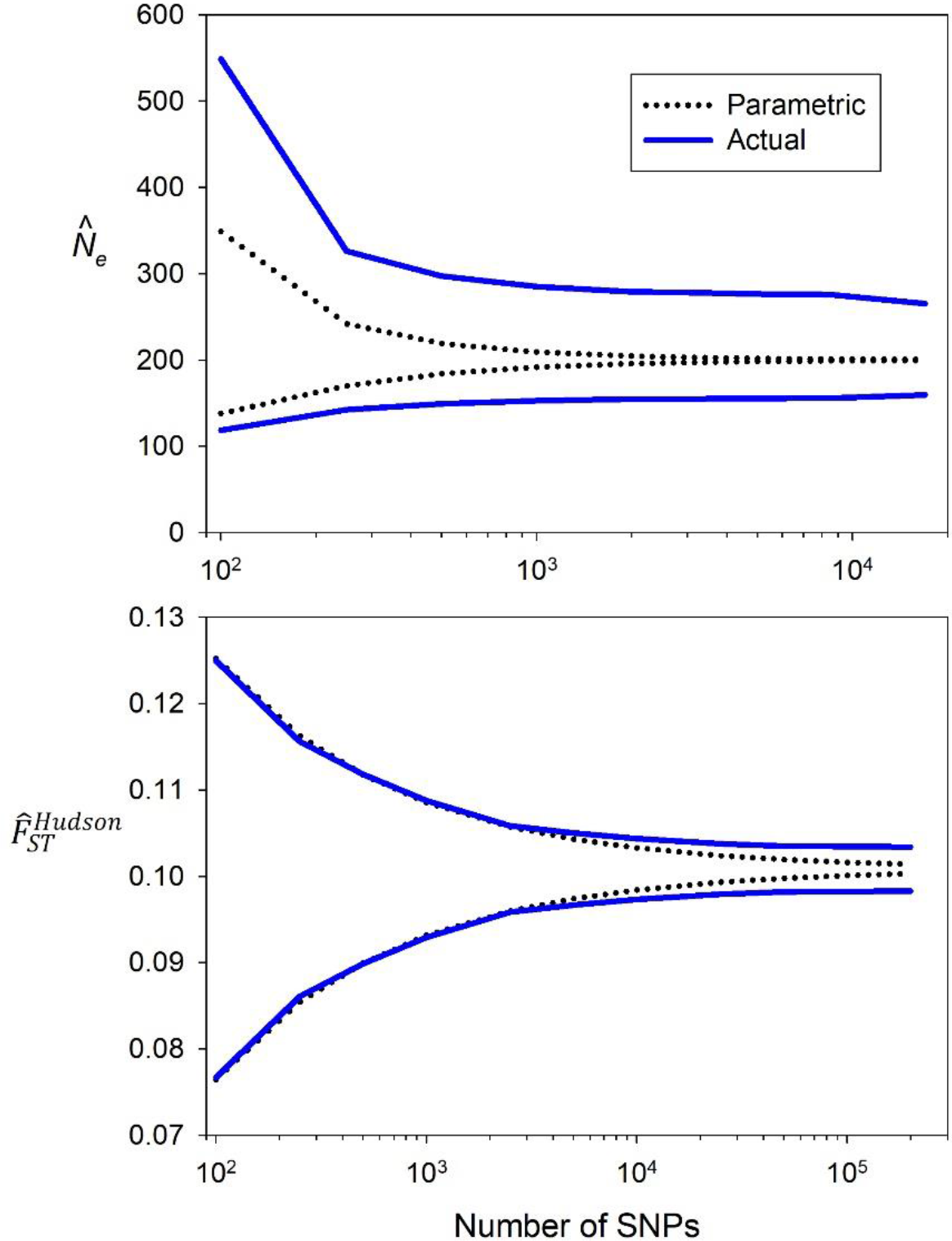
Comparison of parametric and actual 90% confidence intervals for 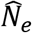 based on LD (top) and 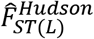 (bottom). Parametric CIs use the nominal degrees of freedom (*L* = the number of diallelic (SNP) loci for 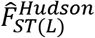; *n* = *L*(*L*-1)/2 for LD); actual CIs use the effective degrees of freedom calculated in this study (*L*’ and *n*’). Results are for simulations with *N*_*e*_ = 200, *C* = 16, and *S* = 50. Note the different X-axis scales in the two panels.

Our results provide a basis for users to develop robust CIs for their own datasets. The best model to predict *n*’ included several sets of covariates for each parameter in the Michaelis-Menten asymptotic function (*p,q,r*). Parameter estimates, standard errors, and more details are included in Supporting Information and Tables S3 and S5. When fit to the original data, the correlation between log(predicted *n*’) and log(true *n*’) was 0.997 (Figure S11). When we evaluated performance of CIs for 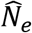 using Equation S5 based on these predicted *n*’ values, the fraction of 90% CIs that contained the true *N*_*e*_ was close to the expected 0.9 (data not shown). But a typical user will know covariate values only for the numbers of individuals and loci in their samples and will have to estimate *N*_*e*_ (from the genetic data, or elsewhere) and perhaps *C* (e.g., from a related species), and finally estimate *n*’ based on our modeling results. Accounting for these additional sources of uncertainty reduced CI performance only slightly, such that overall coverage was 90% for *S*=100, 88% for *S*=50, and 87% for *S*=25 (Table 4).

**Table 4.** Effects of *N*_*e*_, number of chromosomes (*C*), and number of individuals sampled (*S*) on coverage of confidence intervals (CIs) around mean *r*^2^. Results are shown for CIs based on the Jones et al. (2016) jackknife method and effective *df* (*n*’) estimated using the modeling results from this study, which required estimating *N*_*e*_ and *C*. Shown are the percentages of 1024 replicate samples whose 90% CIs included the true value (“In”), were entirely above the true value (“Above”), or were entirely below (“Below”). Each cell represents results averaged over simulations with data for 500, 1000, and 5000 SNPs, and the bottom set of rows averages results across all scenarios, by sample size and method. Results shown used data for pairs of loci on different chromosomes.

| $N_e$ | $C$ | $S$ | Jackknife | | | This study | | |
| --- | --- | --- | --- | --- | --- | --- | --- | --- |
|  |  |  | %Above | %Below | %In | %Above | %Below | %In |
| 200 | 4 | 25 | 9.4 | 1.7 | 88.9 | 12.7 | 1.8 | 85.5 |
| 200 | 4 | 50 | 7.1 | 2.3 | 90.6 | 8 | 3.8 | 88.2 |
| 200 | 4 | 100 | 4.9 | 2.5 | 92.7 | 4.6 | 3.6 | 91.8 |
| 200 | 16 | 25 | 9.5 | 1.3 | 89.1 | 9.4 | 2.4 | 88.2 |
| 200 | 16 | 50 | 5.8 | 1.6 | 92.6 | 7.1 | 5.0 | 87.9 |
| 200 | 16 | 100 | 3.5 | 1.7 | 94.8 | 4.7 | 4.4 | 90.9 |
| 800 | 4 | 25 | 14.8 | 0.6 | 84.6 | 11.7 | 0.8 | 87.5 |
| 800 | 4 | 50 | 8.4 | 1.2 | 90.4 | 10.1 | 1.5 | 88.4 |
| 800 | 4 | 100 | 6.1 | 2.7 | 91.2 | 7.4 | 4.5 | 88.1 |
| 800 | 16 | 25 | 17.1 | 0.5 | 82.4 | 11.4 | 1.2 | 87.5 |
| 800 | 16 | 50 | 8.9 | 1.0 | 90.1 | 9.7 | 2.9 | 87.4 |
| 800 | 16 | 100 | 5 | 1.9 | 93.1 | 6.1 | 3.7 | 90.1 |
| Means |  | 25 | 12.7 | 1.0 | 86.2 | 11.3 | 1.6 | 87.2 |
|  |  | 50 | 7.5 | 1.5 | 90.9 | 8.7 | 3.3 | 88.0 |
|  |  | 100 | 4.9 | 2.2 | 92.9 | 5.7 | 4.1 | 90.2 |

Performance of the Jones et al. (2016) jackknife method varied with sample size (Table 4), which is not surprising given that this method jackknifes over individuals. For *S*=100 the method produced conservative 90% CIs that included the true *N*_*e*_>93% of the time, indicating that *n*’ was generally underestimated. Overall coverage was close to the expected 90% for *S*=50 but only 87% for *S*=25, indicating a tendency of the jackknife method to overestimate precision for small samples. Across all scenarios, most of the jackknife CIs that did not contain the true value were too high, meaning that the lower bound was larger than true *N*_*e*_, and this effect was much stronger for smaller samples.

Scatterplots of estimated *n*’ vs mean *r*^2^ for individual datasets (Figure S12) help to explain performance of Jones’s jackknife method. First, 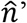 varied considerably across replicates, spanning one order of magnitude for *S*=100 and more than two orders for *S*=25. Second, for all scenarios we found a strong, negative correlation between 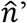 and mean *r*^2^. Datasets with below-average mean *r*^2^ consistently were estimated to have a relatively high *n*’ and hence relatively narrow CIs. Relatively small *r*^2^ translates into a relatively large estimate of *N*_*e*_, around which the CI was relatively narrow. In combination, these factors produce an excess of large 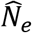 estimates whose lower CI bound is larger than true *N*_*e*_, and this effect is exacerbated for small *S*. Because *n*’ is an increasing function of *N*_*e*_, estimates of *n*’ based on modeling results in this study also show a negative correlation between 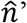 and mean *r*^2^, but the range of 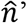 for the current study was about half that of the Jones jackknife method (Figure S12).

### 3.2 *F*_*ST*_

An initial sensitivity analysis (Figures S13-S14 and Supporting Information) showed that Nei’s weighted 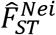 performed better than the unweighted version, and ascertainment method 3 performed better than the other options. Therefore, results that follow apply to 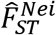 and 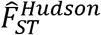 with a MAF cutoff of 0.05.

With no pseudoreplication, variance of 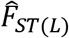 should be inversely proportional to *L*; the actual rate of decline in 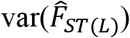 as the number of SNPs increases is shown in Figure 6. For a scenario with *N*_*e*_=200, *C*=4, and *S*=25, the decline is approximately log-linear up to *L*≈2×10^3^, after which point addition of more loci does little to further reduce the variance. For a scenario with larger *N*_*e*_, *S*, and genome size, the rate of decline in 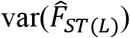 does not start to plateau until the number of loci is an order of magnitude larger (Figure S15). Notably, in both scenarios 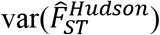 declines at the same rate as 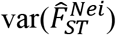 (Figure 6).

**Figure 6.**
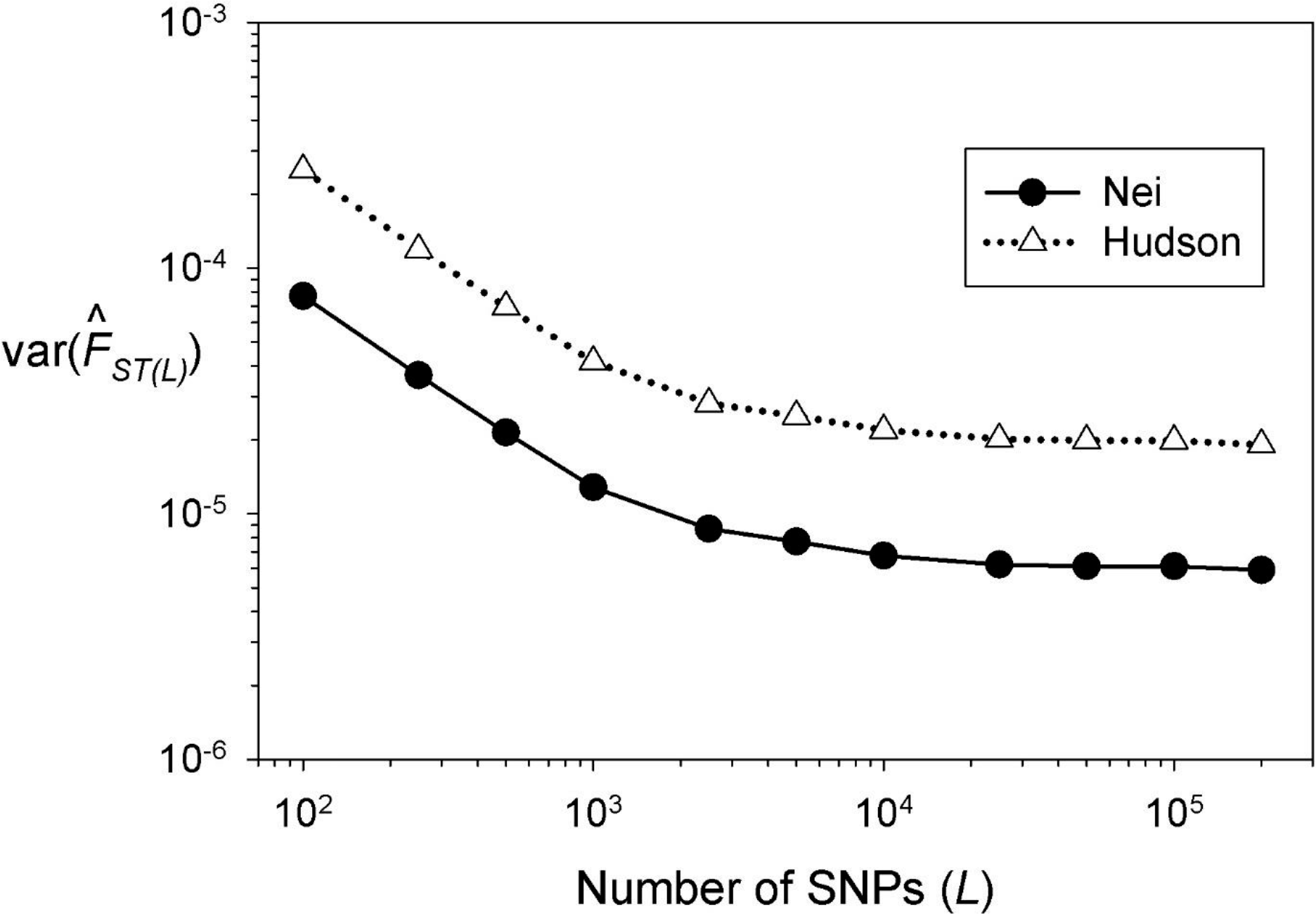
Rate of decline in the variance of multilocus 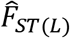 as more diallelic loci (SNPs, *L*) were used in the analysis. Results are for simulations with *N*_*e*_ = 200, *C* = 4, and *S* = 25 and are shown for the estimators of 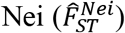 and 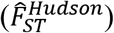. Figure S15 shows comparable results for another scenario with different values of *N*_*e*_, *C*, and *S.*

For a given sample size, mean values of the Nei and Hudson estimators are perfectly correlated with a common slope and sample-size specific intercepts (Figure S16); the slopes and intercepts, however, vary with mean 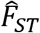. This linear relationship explains why the rate of decline in 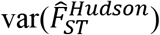 with increasing numbers of loci is nearly identical to that for 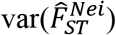.

#### 3.2.1 Effective degrees of freedom

Table S4 gives our estimates of *L*’ for *F*_*ST*_ for every scenario we simulated. As with LD, we found that *L*’ for *F*_*ST*_ depends strongly on both *N*_*e*_ and genome size (Figure 7). In contrast to the two-locus results, however, we found a clear *F*_*ST*_ asymptote for *L*’ only for smaller values of *N*_*e*_ and *C*; for larger values, *L*’ was still increasing after bringing 200K loci into the analysis. We found a moderate effect on *L*’ of sample size of individuals, which became more pronounced for large numbers of loci (Figure S17).

**Figure 7.**
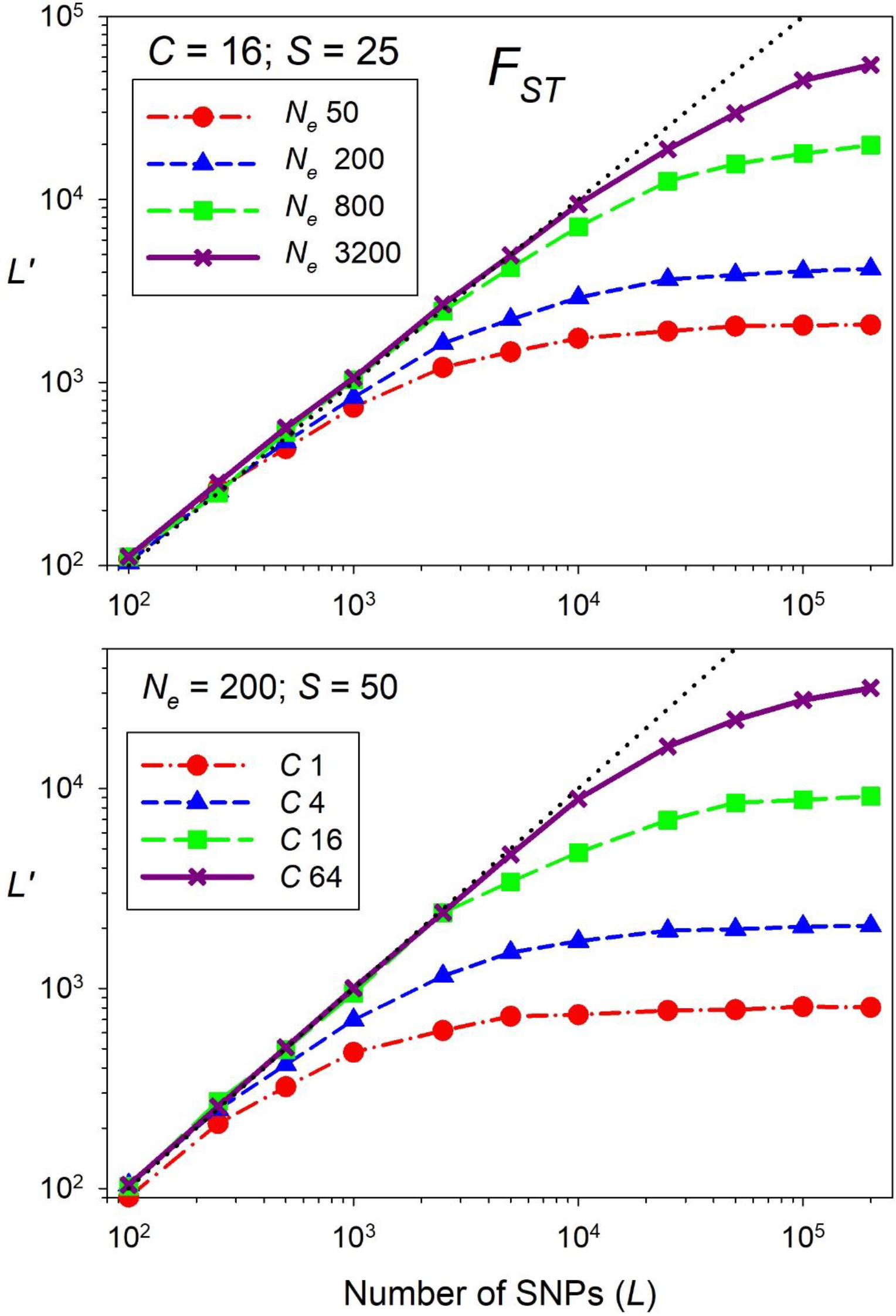
Influence of *N*_*e*_ (top panel, with number of chromosomes, *C,* fixed at 16) and *C* (bottom panel, with *N*_*e*_ fixed at 200) on the effective degrees of freedom (*L*’) for 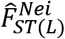 computed between pairs of populations. Black dotted line represents *L*’ = *L* = the number of SNPs.

For a given number of loci, *L*’/*L* was considerably higher for *F*_*ST*_ than *n*’/*n* was for LD (Figure S18). Nevertheless, in populations with small *N*_*e*_ and few chromosomes that are assayed for large numbers of loci, the effective *df* associated with mean *F*_*ST*_ can be two orders of magnitude or more smaller than the number of SNPs. The variance components analysis for *F*_*ST*_ (Figure S19) produced a general pattern similar to that for LD (Figure 4), with one important difference. For the same evolutionary scenario (*N*_*e*_=200, *C*=16, *S* = 50), whereas *V*_2_ for LD rapidly diverges from *V*_1_ and starts to plateau after ~500-1000 loci, the divergence comes much later for *F*_*ST*_, and *V*_2_ does not begin to level off until 10^4^–10^5^ loci are included.

Figure 8 provides a detailed look at variation in 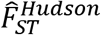 across 48 samples from each of two 2-population pedigrees that produced substantially different mean 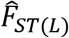 values. Within each pedigree, for each of eight samples of individuals, variation among six replicate samples of *L* loci reflects variance component *V*_1_.

**Figure 8.**
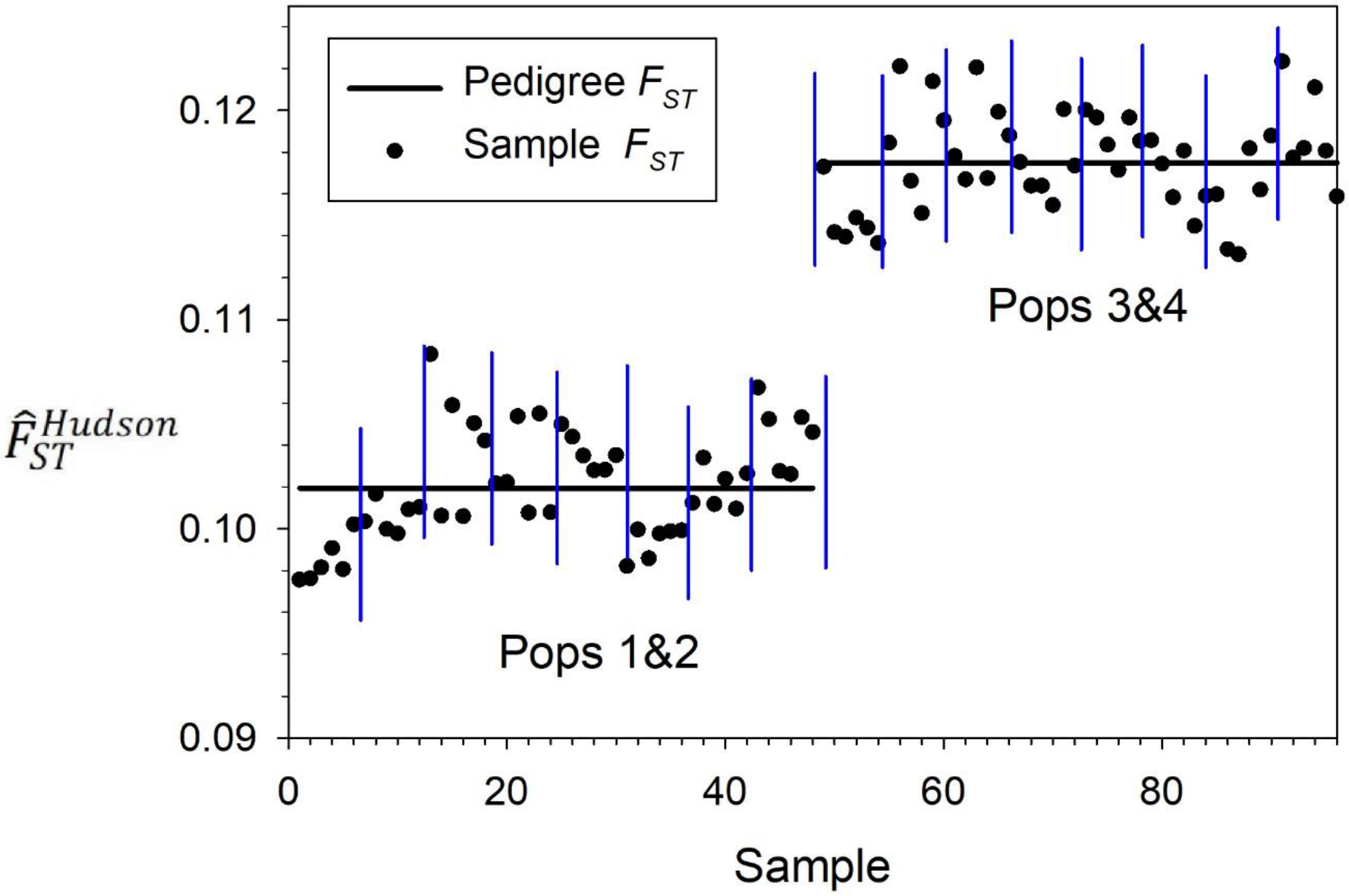
Effects of pedigree on variation in mean 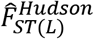. For each of two, 2-population pedigrees, 8 replicate samples (demarcated by vertical lines) were taken of *S* = 100 individuals. These results are for simulations with *N*_*e*_ = 200 and 4 chromosomes. Sampled individuals were drawn hypergeometrically from the *N*_*e*_ individuals in the final generation. For each sample, six mutational replicates generated non-overlapping sets of *L* = 5000 SNP loci that were used to compute mean 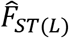. Solid horizontal lines (“Pedigree *F*_*ST*_”) represent mean 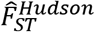 across all 8×6 = 48 replicates within each pedigree. The first set of samples shows results for comparison of daughter populations 1 and 2 and the second set of samples shows results for comparison of daughter populations 3 and 4, all derived from the same ancestral population.

Our estimates of *L*’ for 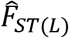 can be used to estimate realistic variances for 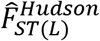 as follows: (1) Given that *L*’ = 2/*ϕ*_*F*_ and 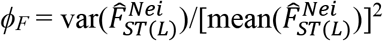, the variance of Nei’s multilocus estimator declines according to 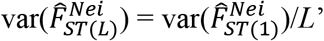, where 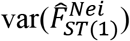 is the variance among single-locus 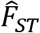 values and 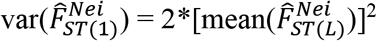. (2) Because 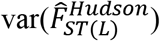 declines at the same rate as 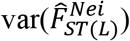, 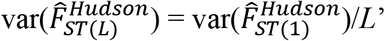, where 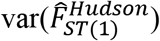 is the variance of single-locus estimates and can be calculated from empirical data.

#### 3.2.2 Confidence intervals

The best model to predict *L*’ for *F*_*ST*_ was similar in form to that found for LD (Supporting Information). When fit to the original data, the correlation between log(predicted *L*’) and log(true *L*’) was >0.99 (Figure S11). CIs based on modeled estimates of 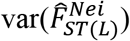 obtained in this study contained the true value 89-95% of the time (mean 91.5%), and of those that did not, roughly equal numbers were too high and too low (Figure 9 and Table 5).

**Figure 9.**
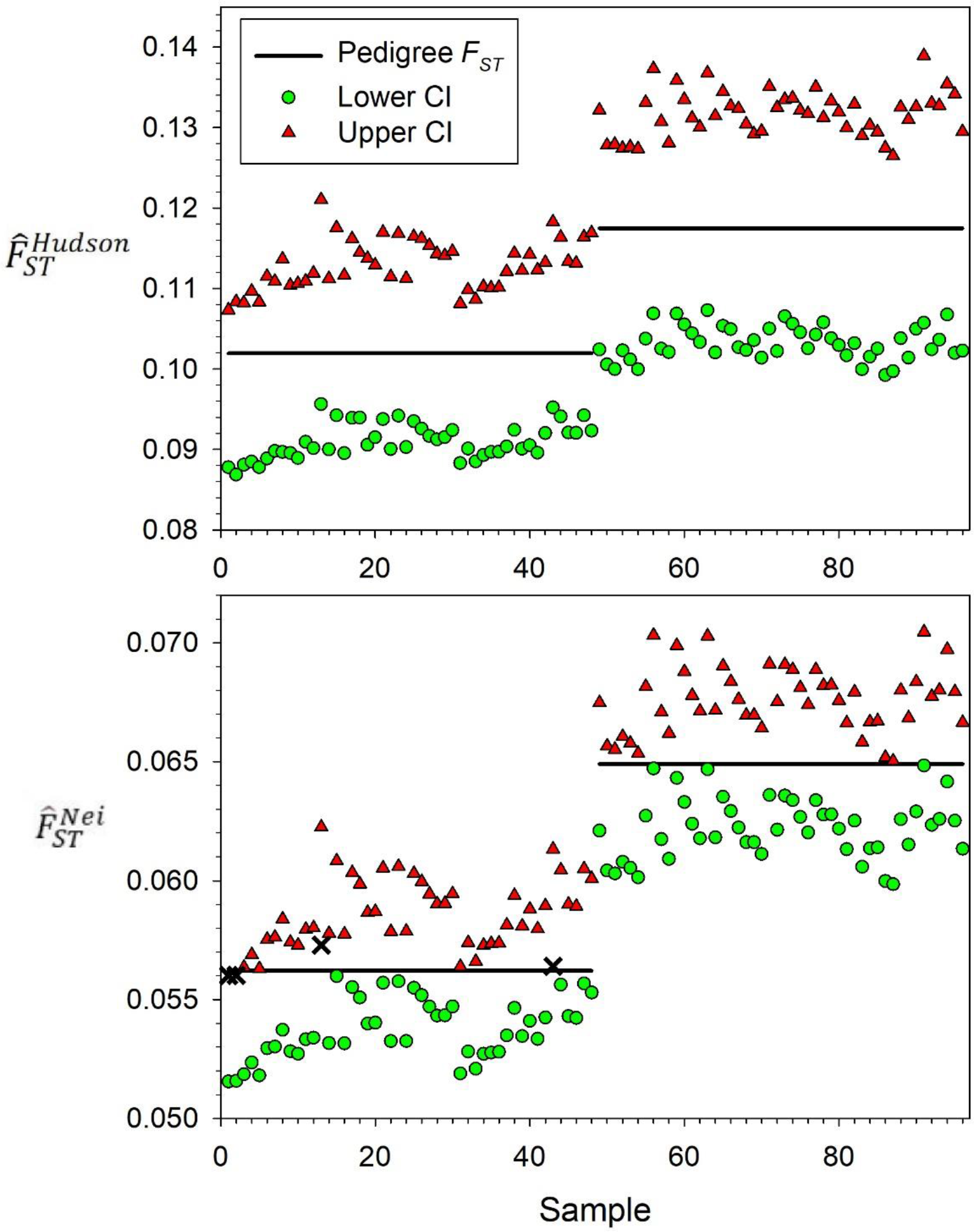
Coverage of 90% confidence intervals (CIs) around *F*_*ST*_ estimators for the population pedigrees and samples shown in Figure 8 (*N*_*e*_ = 200; *C* = 4; *L* = 5000; *S* = 100). Top: CIs generated from block-jackknife estimates of 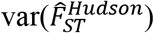. Bottom: CIs generated based on *L*’ for 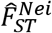 estimated from this study. CI coverage is evaluated with respect to mean 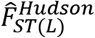 or mean 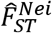 across all replicates within each pedigree (“Pedigree *F*_*ST*_”, horizontal lines). The black X symbols indicate an upper (or lower) bound that was below (or above) the mean pedigree *F*_*ST*_.

**Table 5.** Effects of *N*_*e*_, number of chromosomes (*C*), and number of individuals sampled (*S*) on coverage of confidence intervals (CIs) around 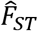. Results are shown for CIs based on block jackknife estimates of 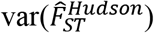 and effective *df*(*L*’) for 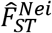 estimated using the modeling results from this study, which required estimating *N*_*e*_ and *C*. Shown are the percentages of 1152 replicate samples whose 90% CIs included the true value (“In”), were entirely above the true value (“Above”), or were entirely below (“Below”). Each cell represents results averaged over simulations with data for 5000, 20000, and 50000 SNPs, and the bottom set of rows averages results across all scenarios, by sample size and method. These results used a block size of 5 Mb. Very similar results were found for block sizes of 1 chromosome and for block-jackknife estimates of 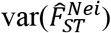 (see Table S7).

| $N_e$ | $C$ | $S$ | Block jackknife | | | This study | | |
| --- | --- | --- | --- | --- | --- | --- | --- | --- |
|  |  |  | %Above | %Below | %In | %Above | %Below | %In |
| 200 | 4 | 25 | 0.7 | 0.6 | 98.7 | 2.4 | 3.9 | 93.7 |
| 200 | 4 | 50 | 0.0 | 0.0 | 100.0 | 2.3 | 2.7 | 94.9 |
| 200 | 4 | 100 | 0.0 | 0.0 | 100.0 | 4.0 | 5.4 | 90.6 |
| 200 | 16 | 25 | 0.2 | 0.8 | 99.0 | 2.4 | 3.8 | 93.8 |
| 200 | 16 | 50 | 0.2 | 0.2 | 99.6 | 2.9 | 4.0 | 93.1 |
| 200 | 16 | 100 | 0.1 | 0.1 | 99.8 | 3.8 | 4.6 | 91.6 |
| 800 | 4 | 25 | 0.5 | 0.3 | 99.2 | 3.0 | 5.6 | 91.3 |
| 800 | 4 | 50 | 0.1 | 0.5 | 99.3 | 4.4 | 6.6 | 89.0 |
| 800 | 4 | 100 | 0.1 | 0.1 | 99.8 | 4.7 | 6.4 | 88.9 |
| 800 | 16 | 25 | 0.7 | 1.2 | 98.1 | 3.8 | 4.8 | 91.4 |
| 800 | 16 | 50 | 0.5 | 1.2 | 98.4 | 3.7 | 5.7 | 90.6 |
| 800 | 16 | 100 | 0.4 | 0.8 | 98.8 | 5.1 | 5.4 | 89.5 |
| Means |  | 25 | 0.5 | 0.7 | 98.7 | 2.9 | 4.5 | 92.5 |
|  |  | 50 | 0.2 | 0.5 | 99.3 | 3.3 | 4.8 | 91.9 |
|  |  | 100 | 0.2 | 0.3 | 99.6 | 4.4 | 5.4 | 90.2 |

Across all scenarios [*N*_*e*_,*C,L,S*], we found three consistent patterns in block jackknife results, and each applied equally to the Nei and Hudson estimators. (1) Block jackknife estimates of variance are positively correlated with mean 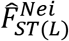 and mean 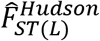 (Figure S20); (2) Virtually all block jackknife estimates exceeded the actual variances we calculated from our simulations (Figure 9 and Table 5); (3) Replicate samples generated using the same parameters produced block jackknife estimates of variance that ranged widely in magnitude (Figure S20), and this variation was greater for the larger block size (one chromosome) and smaller samples of individuals. Because the 5Mb blocks showed less variation among replicates, we focus on those results. Upward bias in the block jackknife estimates of variance produced overly conservative confidence intervals that almost always contained the true values of 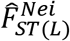 and 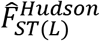 (98-100% coverage for 90% CIs for both Hudson and Nei estimators; Figure 9 and Tables 5 and S7).

R code that allows users to predict *df’* for their own data or other scenarios, for both *F*_*ST*_ and *r*^2^, is available at https://github.com/nwfsc-cb/pseudorep.

## 4 DISCUSSION

The common scenario considered in this paper involves a researcher who has collected data for large numbers of genetic markers in one or a few actual populations. All real populations have a single multigeneration pedigree, and a typical goal is to use genetic methods to draw inferences about evolutionary process that helped shape that population pedigree. Our primary interest is on quantifying uncertainty arising from two sources: sampling genes, and sampling individuals. We do this by simulating many replicate populations and measuring how fast variances of key genetic parameters decline as more loci are used and individuals are sampled.

A substantial complication arises from the fact that the Wright-Fisher reproduction process is inherently stochastic and has many possible realizations. As a consequence, replicate WF populations have different multi-generational pedigrees and different mean values for genetic indices like *r*^2^ and *F*_*ST*_ (Cockerham and Weir 1983; Waples and Faulkner 2009), and averaging across this sort of demographic variance would inflate our estimates of 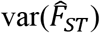 and 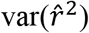 and bias results. To avoid this complication, we used WF reproduction for the forward-in-time component of our simulations, but we took advantage of recently-developed methods (Haller et al. 2019) to generate many replicate samples of genes and individuals drawn from the same population pedigree. This eliminated the demographic component of variance so we could focus on uncertainty associated with sampling of genes and individuals from a single population (or pair of populations). Collectively, our results demonstrate the importance of accounting for differences between pedigrees of the population as a whole and pedigrees of sampled individuals. The variance components analysis showed that for both *r*^2^ and *F*_*ST*_, the primary factor limiting precision in genomics-scale datasets is uncertainty associated with sampling individuals, and this uncertainty cannot be eliminated by sampling arbitrarily large numbers of genes for the same individuals. This is an important consideration for experimental design in genomics-scale datasets, as often it is faster, easier, and cheaper to assay large numbers of genes for a small number of individuals, rather than the reverse.

Effects of pedigrees on statistical inference have been noted previously (e.g., Laurie and Weir 2003; Wakeley et al. 2012, 2016; Ralph 2019). The standard coalescent treats every independent gene as if it were produced on a different pedigree, but in real populations all genes have to percolate through the single, fixed pedigree that captures the genealogical relationships among individuals in the current population and their ancestors, and this pedigree-dependence creates correlations among alleles at different gene loci, even across chromosomes (Bhaskar and Song 2009). This effect is strongest for analyses that are sensitive to pedigrees from the most recent generations, such as relatedness, admixture, population differentiation, and LD (King et al. 2018; Nelson et al. 2020). Furthermore, departures from the standard coalescent model increase when sample size is more than a small fraction of effective size (Bhaskar et al. 2014), a condition commonly encountered in real-world applications.

### 4.1 Linkage Disequilibrium

Our simulations show that, except in populations with large *N*_*e*_ and large genomes, once a few thousand diallelic loci are used to estimate multilocus *r*^2^, adding more loci does little to further reduce 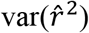. As a consequence, for datasets with 10^4^ or more SNPs, *n*’ can be many orders of magnitude smaller than the number of pairs of loci. Put another way, except when the entire population was sampled, we never estimated *L*’ for LD to be as high as 700 effective loci, even using *L* = 50K SNPs for the largest finite *N*_*e*_ (3200) for the largest genome size (64 chromosomes) that we modeled (Table S2). This in turn means that confidence intervals for 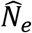 are much wider than they would be if all the pairwise comparisons were independent. The modeling results find significant effects of *N*_*e*_, *C*, *S* and their interactions on *n*’, with *N*_*e*_ having the strongest influence.

The fact that *n*’ depends heavily on *N*_*e*_ even for non-syntenic loci indicates that physical linkage is not the major factor creating lack of independence of pairwise *r*^2^ values; instead, most pseudoreplication arises from overlapping pairs of the same loci in multiple pairwise comparisons. Surprisingly (but conveniently), *n*’ differs very little whether all pairwise comparisons are used or only those on different chromosomes. Restricting comparisons to non-syntenic loci reduces the number of locus pairs (*n*) but simultaneously increases *n*’/*n*, and the two factors effectively offset each other.

It is somewhat ironic that the degree of physical linkage has relatively little effect on pseudoreplication in analyses of LD, but this result can be understood when one considers what happens as more loci are packed into a fixed number of chromosomes. Any new locus will inevitably be linked with many existing loci on whatever chromosome it ends up on, but for every such pairing there are (*C*-1)/*C* new comparisons with existing loci on other chromosomes, and these dilute any effects of linkage. Here we quantify for the first time the large amount of pseudoreplication that occurs because each locus appears in many pairwise comparisons. This overlapping-pairs-of-loci effect is modulated through the population pedigree and is strongest when *N*_*e*_ is small.

Values of *n*’ estimated by the nes et al. Jones et al. (2016) jackknife over individuals were close to, but on average slightly smaller than, overall *n*’ we calculated from simulated data. For replicate datasets simulated using the same fixed parameters, jackknife estimates of *n*’ varied widely and were negatively correlated with mean *r*^2^, which produced relatively narrow CIs when *N*_*e*_ was estimated to be relatively large. Collectively, these features cause results of the jackknife method to be somewhat unpredictable, being on average slightly conservative for larger sample sizes and the opposite for *S*=25, with the latter producing a large excess of CIs that were higher than true *N*_*e*_. Overall coverage of CIs based on *n*’ values estimated by our modeling results was close to the target 90%. However, because of the strong positive correlation between 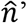 and *N*_*e*_, our modeling results also produce CIs that are relatively narrow when 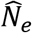 is relatively high, an effect that can be reduced by setting an upper limit to 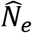 when estimating *n*’.

### 4.2 F_ST_

Consequences of pseudoreplication for precision of 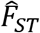 are less dramatic than those for *r*^2^, but nonetheless not trivial. *L*’ approaches an asymptote for relatively small genomes and small *N*_*e*_, but for much larger numbers of loci (~10-20K) than is the case for *r*^2^. For relatively large effective and genome sizes, *L*’ was still increasing after 200K loci, indicating that in those circumstances precision of 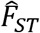 can be enhanced by very large numbers of loci.

Increasing *C* from 1 to 4 increased *L*’ more than any comparable increases in *N*_*e*_. However, most higher organisms have *C≥*4 (Table 1), in which case comparable increases in genome size and effective size produce roughly similar increases in precision. Relatively speaking, increases in sample size have somewhat more effect on *L*’ for 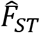 than they do for *n*’ for 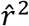. Although the ascertainment and estimation methods we evaluated affect mean values of the index, these differences do not affect the rate at which 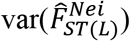 or 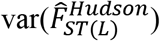 declines with addition of more loci. In related evaluations, we found that changes in the variance of temporal *F* (Nei and Tajima 1981; Jorde and Ryman 2007) parallel those for *F*_*ST*_ (unpublished data). This means that results obtained here can be applied broadly to predict realized precision of temporal *F* statistics and related measures in genomics-scale datasets.

We found that widely-used block jackknife methods consistently overestimate 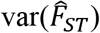, leading to CIs that are much too conservative. In addition, for a given parameter set, the estimated jackknife variance varied several-fold across replicates and was positively correlated with the mean. According to Busing et al. (1999), block size should be large enough to encompass all non-independence, which suggests the appropriate block size should be one chromosome. Although the common block size of 5Mb might be large enough to capture correlations involving sites near the center of the block, this approach arbitrarily divides a continuous system of correlated loci in a way that guarantees that many tightly-linked pairs will be in different blocks. Despite this apparent drawback, the undesirable attributes mentioned above were less extreme for the 5Mb blocks, presumably because they produced more datapoints to analyze. CIs for 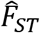 based on *L*’ estimated according to results of this study performed well, even after accounting for uncertainty associated with estimating *N*_*e*_ and *C*.

### 4.3 Experimental Design and Practical Applications

Our simulation and modeling results demonstrate that robust estimates of *df’* can be obtained as a function of numbers of loci and individuals, genome size, and *N*_*e*_. The first two covariates are under control of the investigator, and the third can generally be approximated reasonably well, even for non-model species. Dependence of *df’* on *N*_*e*_ introduces a complication, but with even moderate amounts of genetic data one can obtain a fairly precise estimate of *N*_*e*_ using either single-sample or two-sample (temporal) methods. Even after having to estimate *N*_*e*_ to predict *n*’, LD-based confidence intervals for 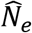 performed at least as well as those obtained using the Jones et al. (2016) jackknife method, and with less variability among replicates (Table 4; Figure S12). Why the block-jackknife method consistently overestimates 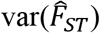 and produces CIs that are too wide is not clear, but it might be related to the fact that blocks with arbitrary boundaries within chromosomes do not capture all dependencies among loci. In any case, using our modeling results to predict *L*’ produced robust CIs for 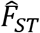. Our results should be particularly useful in planning and experimental design, as expected precision for a wide range of scenarios can be evaluated quickly and easily.

The simulation framework we used, which combines coalescent simulations of the distant past with fast and efficient Wright-Fisher forward simulations of the recent past, provides more realistic results than can be achieved by either process alone (Nelson et al. 2020). Nevertheless, as is inevitable our model required simplifying assumptions, so some caveats are in order. We assumed closed populations and did not evaluate potential consequences of migration for 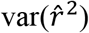 or 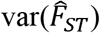. We modeled non-random associations of neutral genes and did not attempt to account for correlations due to selective pressures on linked or unlinked sites, so in that respect our estimates might be considered upper limits to actual *df’*.

We did not explicitly model variation in recombination rate, which is known to be common both across the genome and between sexes (Ritz et al. 2017; Sardell and Kirkpatrick 2020), so our results reflect a generic genome-wide average. Although the genome sizes we simulated (1-64 chromosomes of 50 Mb) encompassed the range of mean values reported for higher organisms (Table 1), all chromosomes we modeled were the same size. Some effects of unequal chromosome length can be accounted for by defining an effective number of chromosomes as 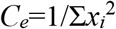, where *x*_*i*_ is the relative length of the *i*^*th*^ chromosome, standardized such that Σ*x*_*i*_=1. *C*_*e*_ is analogous to the effective number of alleles at multiallelic loci. However, *C*_*e*_ only deals with interactions among chromosomes and does not account for different patterns of recombination within chromosomes.

A more general formulation that considers both intra- and inter-chromosomal effects on genetic shuffling was proposed by Veller et al. 2019 PNAS 116:1659, who defined a metric 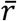, which is “the probability that the alleles at two randomly chosen loci are shuffled in the production of a gamete” (p. 1660). In Veller et al.’s framework, the expected value of 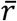 can be expressed as the sum of two terms: an intra-chromosomal term that is the probability that two loci are on the same chromosome and shuffle their alleles; and an inter-chromosomal term that is the probability that two loci are on separate chromosomes and shuffle their alleles.

As shown in Supporting Information, the effective number of chromosomes accounts for the inter-chromosomal effect on 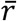, which quantifies effects of independent assortment (Mendel’s Second Law), and the distribution of chromosome sizes also affects the intra-chromosomal effect. For any organism with more than a few chromosomes (mean for vertebrates is 25; Table 1), the inter-chromosomal effect on 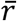 greatly exceeds that of patterns of recombination within chromosomes. Therefore, use of *C*_*e*_ to account for effects of unequal chromosome length should provide a good first-order approximation for the overall amount of genetic shuffling and hence pseudoreplication.

Nevertheless, two additional factors contribute to the intra-chromosomal effect on 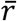: 1) the number of crossovers (COs) on each chromosome, and 2) their locations. All else being equal, more COs lead to more shuffling, and COs near the center of a chromosome lead to more shuffling than crossovers near the ends (Veller et al. 2019). Within chromosomes, our simulations modeled the number of COs as a random Poisson variable, with locations of COs randomly spaced along the chromosome. In Supporting Information, we show (Equations S14-S15) how researchers who are interested in results for species with different patterns of recombination within chromosomes can adjust our results to reflect the desired overall value of 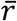 and hence the appropriate level of pseudoreplication.

We modeled discrete generations, and our sampling design assumed that individuals were sampled randomly from the entire adult population. Forward simulations used Wright-Fisher dynamics with a constant number of ideal adults (*N*), so *N*_*e*_≈*N*. We found a qualitative difference in *df’* for samples that included the entire population (*S*=*N*_*e*_), but for real populations (typically with *N*_*e*_<*N*) the relevant criterion is whether all individuals have been sampled (*S*=*N*). Values for *df’* reported in Table S2 for *S*=*N*_*e*_ would provide a robust estimate of expected precision for the special case where it is possible to assay the entire population, thus eliminating the large variance component associated with sampling individuals. Finally, for many species it is most convenient to sample juvenile offspring rather than adults. The variance associated with juvenile samples approximates that for a very small sample of the parents (see Supporting Information). Therefore, an approximate value for *df’* for such samples can be obtained by using the predicted *df’* (*n*’ or *L*’) for small *S*.

## Acknowledgments

We thank Eric Anderson, Nicholas Galwey, Marty Kardos, Peter Ralph, Carl Veller, and John Wakeley for useful discussions and Martin Liermann for suggestions regarding model fitting. An anonymous reviewer provided useful suggestions that improved the manuscript.

## Data Accessibility

This manuscript did not generate new empirical data. Code to conduct the simulations and related analyses is available at https://github.com/nwfsc-cb/pseudorep.

## Author Contributions

RSW conceived the study; RSW and RKW designed the study and simulated and analyzed the data. EW conducted the analyses to model *n*’ as a function of key covariates. RSW wrote the manuscript with input from RKW and EW, and all authors edited the manuscript.

## Notes

https://github.com/nwfsc-cb/pseudorep

### Competing Interest Statement

The authors have declared no competing interest.

### Summary of Updates

Revised to reflect reviewer comments. Computer code has been uploaded to Github

