## Supplemental text and figures for "Pseudoreplication in genomics-scale datasets"

### Supporting Information

Detailed Methods
- Conceptual framework
- Simulations
- Effective degrees of freedom
- Data analysis
- Model fitting
- Confidence intervals
- Effective number of chromosomes and recombination
- Juvenile sampling

References
Figures

See also these Supporting Information Tables in a separate Excel file available on Github at https://github.com/nwfsc-cb/pseudorep

Table S1. Variable transformations used in modeling
Table S2. Effective degrees of freedom (df) for the LD analyses
Table S3. Model fits for the LD analyses
Table S4. Effective degrees of freedom (df) for the $F_{ST}$ analyses
Table S5. Model fits for the $F_{ST}$ analyses
Table S6. Detailed results for block-jackknife estimates of var($F_{ST}$)

#### Detailed Methods

##### Conceptual framework

The Wright-Fisher (WF) model of random mating and reproduction has been a cornerstone of theoretical and applied population genetics for almost a century. The WF process is inherently stochastic, and each realization of WF reproduction produces a population pedigree, which records the genealogical relationships among current and past members of the population. In many population genetic analyses, the population pedigree is considered a nuisance parameter and integrated over to allow broader insights into evolutionary processes. We take a different approach here because our focus is on actual populations, each of which has a single, realized population pedigree (Wakeley et al. 2012; Ralph 2019). Interaction between this population pedigree and mutational processes produces the patterns of genetic variation observed in samples from the population. We are interested applying genetic tools to gain information about the population pedigree and its consequences for applied conservation and management.
Our approach involves simulating many replicate population pedigrees and evaluating how rapidly variances of mean statistics decline when averaged across many thousands of genes (or many millions of pairwise comparisons of genes).

It is essential to carefully consider the processes of replication and sampling, because these variances become very small for large numbers of loci, and any extraneous source of variation can profoundly affect results. In particular, under the WF model, realized effective population size ($N_e$*) varies randomly each generation around the nominal $N_e = N$ ideal individuals, with variance ≈ $N/2$ (Waples and Faulkner 2009). Initially we thought it would be sufficient to control the single-generation population pedigree, which can be accomplished by ensuring that realized variance in reproductive success (and hence realized $N_e$*) is constant each generation. We expected that this would provide two methods of replication that should be equivalent: computing the variance of mean $r^2$ (or mean $F_{ST}$): 1) in samples taken from replicate populations with controlled $N_e$*; or 2) in replicate sets of loci assayed on the same individuals.

However, it turns out that this approach is problematical for two reasons. First, simply eliminating variation in single-generation realized $N_e$* still allows spurious variation among replicate populations because the multi-generation pedigree was not controlled. Second, genetic characteristics of any set of individuals sampled from a population will be influenced by the pedigree peculiar to those individuals. With replication across more genes, estimates of genetic indices will converge on parametric values that are determined by the pedigree of the sampled individuals. Often, however, one wants to draw inference about the population as a whole, and uncertainty related to that cannot be eliminated by even arbitrarily extensive replication across a subset of individuals (Wakeley et al. 2012; King et al. 2018; see also Laurie and Weir 2003; Bhaskar and Song 2009 for discussion of a related issue involving human forensics testing).

To address the first issue, we used a new simulation method (described below) that allowed us to precisely control the multi-generation pedigree of each population. To address the second issue, we modified our experimental design to explicitly account for two sources of uncertainty: sampling a subset ($S$) of individuals from the total population size ($N$), and sampling a subset of loci from the entire genome. As the maximum number of loci we modeled ($2\text{x}10^5$ for $F_{ST}$) was many orders of magnitude smaller than the average genome size in multicellular eukaryotes (~$1.5\text{x}10^9$ bp; Li et al. 2011), we modeled sampling non-overlapping sets of gene loci. In contrast, because it might be possible to sample a substantial fraction (perhaps even all) of the individuals in a population, our replicate samples of individuals were drawn hypergeometrically from the same $N_e = N$ total individuals in the final generation of each simulation replicate. Each of four (potentially partially-overlapping) samples of individuals was analyzed for four non-overlapping sets of loci, creating a 4x4 matrix of mean $r^2$ or $F_{ST}$ values (Table 1). Data from the rows and columns, respectively, were used to calculate variance components $V_1$ (same individuals, different loci) and $V_2$ (same loci, different individuals). Data from the cells on the diagonal (different samples of individuals, different sets of loci) were used to calculate the effective degrees of freedom ($n$' or $L$') as described below.

**Simulations**

SLiM (Messer et al. 2013) and msprime (Kelleher et al. 2016) were used to simulate genetic variation. For each evolutionary scenario, SLiM (versions 3.2.1-3.3.1) was used to simulate 4 ancestral diploid populations of $N_e = N$ ideal individuals for $10N_e$ generations under WF reproduction with separate sexes. Then, each ancestral population split into 4 daughter populations of size $N_e$, which evolved independently under isolation for $t = 0.2N_e$ additional generations. Under isolation, $E(G_{ST}) \approx (s\text{-}1)T/(s\text{-}T)$ (Nei and Chakravarti 1977), where $G_{ST}$ is Nei's (1973) index that is equivalent to Wright's $F_{ST}$ for two alleles, $s$ is the number of subpopulations, and $T = t/(2N_e)$. Here we follow the common practice of computing $F_{ST}$ for pairs of populations, so $s = 2$ and $E(G_{ST}) \approx T/(2\text{-}T) = t/(4N_e - t)$. Setting $E(G_{ST}) = 0.05$ leads to $t \approx 0.2N_e$. This resulted in divergence times of $t$ = 10-640 generations for scenarios with $N_e$ = 50-3200.

Outputs of the forward simulations were saved as tree sequence files (Kelleher et al. 2018) that fully describe the ancestry of each individual. These tree sequence files were "recapitated" using msprime (version 0.7.1), which simulated the distant past to ensure that all individuals fully coalesced. msprime was also used to add mutations to the simulated genealogies. It uses an infinite sites mutation model, so variable sites have only two alleles. The mutation rate used in the simulations was $10^{-8}$, and this was increased as needed to ensure the specified number of variable sites. We did not simulate genotyping errors or missing data. Scripts used for the simulation are available on Github (https://github.com/nwfsc-cb/pseudorep).

Although unlinked loci and infinite $N_e$ are not achievable in any real population, these scenarios are easy to model in a computer and they provide insights into asymptotic behavior as $N_e$ and $C$ become very large. Unlinked loci don't require a coalescent phase to reach linkage equilibrium—a few generations of forward simulations are sufficient. To model unlinked loci, we used WF reproduction and the same ranges of sample sizes and $N_e$ as the SLiM simulations. Mutational replicates were not generated, so replication within each population followed the scheme in Figure 1. Because no genetic drift occurs in a population of infinite size, for the single-locus analyses we simply sampled from the same parametric allele frequencies twice. For the LD analyses, we drew multilocus genotypes randomly and independently from a vector of parametric allele frequencies; this modeled sampling progeny produced by an infinite number of parents.

**Effective degrees of freedom**.

We calculated the effective degrees of freedom (df) based on the observed variances of $\hat{F}_{ST(L)}$ and $\hat{r}^2{}_{(L)}$ across replicate datasets, using properties of the chi-square distribution. Consider a random variable $X_i$, and consider the replicate mean values $\bar{X}_q$ that are each calculated over $q$ independent $X_i$ values. The chi-square distribution has the property that, if $\bar{X}_q$ is distributed as chi square with $q$ degrees of freedom, $E(\bar{X}_q) = q$ and $var(\bar{X}_q) = 2q$. It follows that, for a chi-square variate with $q$ df, this relationship holds:

$\phi = var(\bar{X}_q)/E^2(\bar{X}_q) = 2q/q^2 = 2/q$; and (S1)

$var(\bar{X}_q) = 2E^2(\bar{X}_q)/q$. (S2)

Thus, for a chi-squared variate the variance of the mean is inversely proportional to the degrees of freedom, which is the number of independent elements summed over to produce the mean. Rearranging Eq (S2) produces

$q = 2/\phi = 2E^2(\bar{X}_q)/var(\bar{X}_q)$. (S3)

If the $X_i$ are not independent, then $var(\bar{X}_n)$ will be higher than expected for a chi-square variate with $q$ df. In that event, we can use empirical values of $mean(\bar{X}_q)$ and $var(\bar{X}_q)$ in Eq S3 to calculate an effective degrees of freedom, $q'$. The effective degrees of freedom is the number of independent values of $X_i$ that, if averaged over, would produce the same $var(\bar{X}_q)$ as the observed variance.

The variance of $F_{ST}$ across loci has long been of interest to those interested in identifying "outlier" loci that might be under selection. Lewontin and Krakauer (1973) first proposed the metric $k = Var(\hat{F}_{ST})/\ E^2(\hat{F}_{ST})$, which is the squared coefficient of variation of $\hat{F}_{ST}$ across loci, as a way to identify unusually large single-locus $F_{ST}$ values that might provide evidence for selection. Although $k$ is somewhat sensitive to the allele frequency distribution, Lewontin and Krakauer showed with simulations that $k$ is approximately 2 when $F_{ST}$ is computed over 2 populations and the distribution of allele frequencies is binomial, as expected in early stages of population divergence (Robertson 1975). Some additional factors related to population structure and population history can inflate the variance of $\hat{F}_{ST}$ across loci (Nei and Maruyama 1975; Ewens and Feldman 1976), and addressing these issues has been a major focus of attempts to use this general approach to identify loci under selection (Beaumont and Nichols 1996; Anatao et al. 2008; Narum and Hess 2011) and more recently using genome scans (Lotterhos 2019; Booker et al. 2020). In our experimental design, however, accounting for different population pedigrees deals with the hierarchical population structure issue, and there are no other sources of heterogeneity that could be expected to inflate Var($\hat{F}_{ST}$). Therefore, Lewontin and Krakauer's *k* parameter, which we call $\phi_F$, is a useful metric of pseudoreplication of $F_{ST}$ for our experimental design. The L-K results suggest that, if all *L* diallelic loci are completely independent, the following relationship should hold: $\phi_F = 2/L$, where *L* is the number of independent datapoints. Using extensive simulations, Nei and Tajima (1981), Pollak (1983) and Waples (1989) showed that the same is true for the closely-related temporal *F*. Therefore, we computed the effective degrees of freedom from the empirical variance of $\hat{F}_{ST(L)}$ as $L' = 2/\phi_F$. The value *L'* can be interpreted as the number of independent data points that would be expected to produce the observed variance (Cox 1984; Giesbrecht 2006). The ratio $L'/L$ therefore provides an index of how much pseudoreplication has reduced precision of the estimator. An analogous approach was used with LD based on the actual and effective numbers of pairs of loci, *n* and *n*'.

#### Data analysis

**LD.** For each pair of loci, the sample estimate of the squared correlation coefficient ($\hat{r}^2$) was computed using the Pearson product-moment correlation of alleles at different loci, which produces a result identical to that of the composite Burrows method (Weir 1996; Gao et al. 2008). Across multiple pairs of loci, the unweighted mean ($\hat{r}^2{}_{(L)}$) was used as an overall sample estimate of $r^2$. To estimate $N_e$ using the LD method, we first multiplied $\hat{r}^2{}_{(L)}$ by the factor $[S/(S-1)]^2$ to reflect the bias correction suggested by Weir (1979) that is implemented in LDNe (Waples and Do 2008). Then, $N_e$ was estimated using the sampling adjustments in Table 2 of Waples (2006); with equal sample sizes across loci, this produces the same estimate reported by LDNe and NeEstimator (Do et al. 2014).

***F*$_{ST}$.** Each ancestral population produced 4 daughter populations, which allowed 6 population pairs for estimating $F_{ST}$. The daughter populations have different pedigrees, so (for example) in computing *L*' for the comparison of daughter populations 1 and 2, we computed $\phi_F$ using all 8 mutational replicates for these two populations, then averaged mean $\phi_F$ values across pairs of daughter populations and ancestral populations to obtain an overall mean $\phi_F$, which was converted to *L*'.

In an initial sensitivity analysis, we evaluated different methods of estimating $F_{ST}$, under conditions (relatively few loci and a large number of chromosomes) where effects of linkage should be small and $L'/L$ should be close to 1. Results showed that weighted $\hat{F}_{ST}^{Nei}$ performed as expected, whereas unweighted $\hat{F}_{ST}^{Nei}$ produced estimated *L*' values >> than the number of loci (Figure S13, top). This is consistent with previous reports that show unweighted versions of *F* statistics are more biased but have a smaller variance (Jorde and Ryman 2007; Bhatia et al. 2013) and can produce values of $\phi_F$ <2 (Nei and Tajima 1981; Waples 1989). Of the three ascertainment schemes evaluated for weighted $\hat{F}_{ST}^{Nei}$, no ascertainment and ascertainment in the ancestral population produced estimates of $L' \ll L$, indicating an elevated variance (Figure S13, bottom). Accordingly, results that follow apply to weighted $\hat{F}_{ST}^{Nei}$ and $\hat{F}_{ST}^{Hudson}$ under scheme 3, which applied a MAF cutoff of 0.05.

**Model fitting**

In this section, we describe additional details of the model fitting procedure for the LD and $F_{ST}$ analyses.

**LD Analyses.** In addition to the 3-parameter asymptotic curve described in the main text, we considered multiple versions of the Michaelis–Menten (or Beverton-Holt) equation modified to include covariates. The general form of these functions was

$$f(x) = \theta_1 \cdot z_1 + \frac{(\theta_2 \cdot x + \theta_3 \cdot z_2)}{(\theta_4 + x + \theta_5 \cdot z_3)}, \quad \text{(S4)}$$

where the values $z_1$, $z_2$, and $z_3$ represent optional covariate data, and the parameter vector $\theta$ represents optional parameters (when $\theta_1$, $\theta_3$, and $\theta_5$ = 0, this equation takes the form of the Michaelis–Menten model without covariates. For LD, the number of datapoints (pairs of loci) was $n = L(L\text{-}1)/2$. We explored a wide range of models, using raw covariates (*n*, *S*, *L*, *C*, *Ne*) as well as transformed versions of these described in Table S1.

To allow more flexibility in the shape of the asymptote, a third version of models considered were generalized additive models (GAMs) with shape constraints, using the mgcv (Wood 2004) and scam (Pya 2020) packages in R (R Core Team 2020). Because of a potentially skewed response distribution of $n'$, we considered models either with $n'$ or log($n'$) as the dependent variable. The asymptotic models described in the main text and above were fit using using both sum-of-squares and maximum likelihood techniques (using the bbmle package; Bolker and R Core Team 2020), while the GAM models were fit only using maximum likelihood.

For models described in the main text, we implemented a 2-stage procedure for maximizing the likelihood. First, we generated a smaller dataset representing each combination of $N_e$, *C*, and *S* (for any set of these values, our simulated data included a range of loci). Optimal values of *p*, *q*, and *r* were found on this smaller dataset using sum of squares, and then the coefficients *p*, *q*, and *r* were treated as response variables in regressions against the predictors described in Table S1. Confidence intervals on $n'$ were generated with a Monte Carlo approach, where uncertainty in the 3 sets of regression parameters was simulated over 10000 iterations, and propagated forward via the asymptotic equation as a function of *p*, *q*, and *r* described in the main text.

Using sum of squares and Equation S4, our best fit model for predicting log($n'$) included an asymptote in log($n$),

$$E[\log(n')] = \theta_1 \cdot \left(\frac{Loci}{C}\right) + \frac{(\theta_2 \cdot \log(n) + \theta_3 \cdot \log(Ne))}{(\theta_4 + \log(n))}, \quad \text{(S5)}$$

where the parameters and standard errors (calculated from the Hessian matrix) were estimated to be $\theta_1$ = -0.0001 (SE = 0.0001), $\theta_2$ = 36.6577 (SE = 6.1913), $\theta_3$ = 15.8487 (SE = 1.7682), $\theta_4$ = 30.2171 (SE = 6.0092). For this model, the log-log correlation between predicted and true *n*' was 0.977.

We found a better overall fit using the two-step version of the Michaelis-Menten model described above; our best model (AIC = -571.6690) included several covariates for each of the 3 parameters. In step 1, for each scenario defined by combinations of the covariates *Ne, C*, and *S*, we fit the 3 parameters (*p*, *q*, and *r*) to the set of *n*' values for different numbers of loci. In step 2, we found the best linear combinations of the covariates to predict *p*, *q,* and *r.* The best model included four sets of covariates for each parameter. In this model, log10($n'$) is used as the response. The form of these relationships was

$$p = b_1 + b_2 log(C) + b_3 log(Ne) + b_4 \log(S) + b_5 SNe + b_6 log(C) log(Ne)$$

$$q = b_7 + b_8 log(C) + b_9 log(Ne) + b_{10} log(S) + b_{11} SNe + b_{12} \left(\frac{1}{C}\right) + b_{13} log(C) log(Ne) + b_{14} log(C) log(S)$$

$$r = b_{15} + b_{16} log(C) + b_{17} log(Ne) + b_{18} \log(S) + b_{19} log(C) log(Ne) + b_{20} log(S) log(Ne) + b_{21} log(S) log(C),$$

with coefficients shown in Table S3. When fit to the original data, the log-log correlation between predicted and true $n'$ was 0.997 (Figure S11).

***$F_{ST}$***. For $F_{ST}$ the number of datapoints is the number of diallelic loci, $L$, so the effective df = $L'$. We adopted the same overall framework in modeling $L'$ for $F_{ST}$, with the best model taking the form

$$p = b_1 + b_2 log(C) + b_3 log(Ne) + b_4 log(C) log(Ne)$$

$$q = b_5 + b_6 log(C) + b_7 log(Ne) + b_8 log(S) + b_9 SNe + b_{10} log(C) log(Ne) + b_{11} log(Ne) log(S) + b_{12} log(C) log(S)$$

$$r = b_{13} + b_{14} log(C) + b_{15} log(Ne) + b_{16} log(S) + b_{17} log(C) log(Ne) + b_{18} log(Ne) log(S) + b_{19} log(C) log(S),$$

with coefficients shown in Table S5. When fit to the original data, the log-log correlation between predicted and true $L'$ was > 0.99 (Figure S11).

**Confidence intervals**

To evaluate accuracy of our estimates of the degree of pseudoreplication, for selected scenarios we generated many (>1000) new replicate samples, and for each sample we calculated $\hat{F}_{ST(L)}$ and $\hat{r}^2{}_{(L)}$ averaged over data for 100–5000 SNPs ($\hat{r}^2$) or 5000–50000 SNPs ($\hat{F}_{ST}$). For the LD analyses, for each of the 16 population pedigrees we drew 8 new samples from each of the 8 mutational replicates, which produced a total of 16x8x8 = 1024 new samples. For $F_{ST}$, for each of the 24 two-population pedigrees, we took 8 new samples from each of the 6 mutational replicates, producing 1152 new samples. For all sample × mutation replicates derived from the same pedigree, the overall mean $\hat{F}_{ST(L)}$ or $\hat{r}^2{}_{(L)}$ was taken as the parametric value for that pedigree. Confidence intervals (CIs) were then applied to the point estimate for each replicate, and the fraction of CIs that included the parametric value for that pedigree was recorded as the coverage. This ensured that the coverage data reflected only precision and not systematic biases in the estimators or random variation across population pedigrees.

CIs for point estimates of $\hat{F}_{ST(L)}$ and $\hat{r}^2{}_{(L)}$ were generated using standard statistical theory. For the LD analyses, simple functions of mean $\hat{r}^2$ are distributed as chi square (Hill 1981). This can be used to set CIs for $\hat{r}^2$ using an analogue of Equation S7 (Waples 2006):

$$(1\text{-}\alpha) \text{ CI for } \overline{\hat{r}^2} = \left[\frac{n'\overline{\hat{r}^2}}{X^2{}_{\alpha/2[n']}}, \frac{n'\overline{\hat{r}^2}}{X^2{}_{1-\alpha/2[n']}}\right], \quad \text{(S6)}$$

where $n'$ is the effective df = effective number of pairs of loci. CIs for $\hat{N}_e$ are then generated by using the critical $\hat{r}^2$ values to estimate $N_e$ as in Waples (2006).

For the single-locus analyses, CIs were calculated as

$$\text{CI}(\hat{F}_{ST(L)}) = [\hat{F}_{ST(L)} \text{-} z\sigma_{\bar{\hat{F}}} , \hat{F}_{ST(L)} + z\sigma_{\bar{\hat{F}}}], \quad \text{(S7)}$$

where $z$ is the standard normal deviate and $\sigma_{\bar{\hat{F}}}$ is the standard deviation of $\hat{F}_{ST(L)}$. We evaluated two-tailed coverage for 90% CIs, so we used $z = 1.645$. Alternatively, CIs for Nei's $\hat{F}_{ST(L)}$ can be set based on the chi square distribution (after Waples 1989):

$$(1\text{-}\alpha)\text{ CI for }\hat{F}_{ST(L)} = \left[\frac{L'\hat{F}_{ST}}{X^2{}_{\alpha/2[L']}}, \frac{L'\hat{F}_{ST}}{X^2{}_{1-\alpha/2[L']}}\right], \quad \text{(S8)}$$

where $\chi^2{}_{\alpha/2[L']}$ is the α/2 point of the chi-square distribution with $L'$ degrees of freedom. Although the distribution of $\hat{F}_{ST}$ is skewed for small numbers of loci (Lewontin and Krakauer 1973), the normal approximation is very good for the large numbers of SNPs considered here, in which case Equations S6 and S7 produce essentially identical results.

Width of the CIs was calculated in two ways. First, for each scenario we modeled the expected value of effective df as a function of the covariates $N_e$, $C$, $S$, $L$, as described in the previous section. $S$ and $L$ are observed from the data; we assumed the chromosome number ($C$) was known within a random error term (with 25% CV), and $N_e$ was estimated from observed $\hat{r}^2$ according to Waples (2006). For the single-locus analyses, $\hat{N}_e$ was computed as the harmonic mean of the estimates for the two populations. Because $\hat{N}_e$ is skewed high and can be arbitrarily large, we set an upper limit to $\hat{N}_e$ of $10^4$ for $F_{ST}$ and 2000 for LD. These evaluations modeled a scenario in which a user has one or more empirical datasets for which he/she wants to set confidence limits, and can obtain an estimate of effective df from our modeling results and the empirical data.

Second, for comparative purposes, we also calculated CIs for each dataset using a published jackknife procedure. To account for lack of independence due to linkage in single-locus genetic metrics, investigators having dense genomics data often use a block jackknife (Efron 1982; Busing 1999), in which blocks of loci (rather than individual loci) are sequentially omitted from computation of the index of interest. We calculated mean $\hat{F}_{ST(L)}$ for each of $M$ blocks, and the square root of $M$ times the variance of these $\hat{F}_{ST(L)}$ values was used as a standard error to place normal-distribution confidence limits around $\hat{F}_{ST(L)}$, using Equation S6 (cf. Reich et al. 2009; Durand et al. 2011). For these analyses, we retained the genomic structure of the data, with loci arranged sequentially along each chromosome. The blocks should be larger than the regions affected by lack of independence (Busing 1999). Because linkage is continuous across a chromosome, this implies that the block size should be one chromosome. However, researchers using this method with genomics data for humans or model species typically target individual chromosomes or smaller regions, and 5 Mb is a typical block size used (e.g., Reich et al. 2009). Therefore, we evaluated block sizes of one chromosome and 5 Mb.

The block jackknife is not suitable for obtaining a variance of $\hat{r}^2{}_{(L)}$ because the units of information involve pairs of loci, and there is no way to sequentially remove non-overlapping sets of loci without affecting many overlapping pairs of loci. However, Jones et al. (2016) developed a standard jackknife for the LD method that sequentially omits individuals rather than loci. For each sample, we used the Jones et al. method, including their correction factor $0.84^2$ for the estimated jackknife variance, to estimate the variance of $\hat{r}^2{}_{(L)}$ and from this the estimated $n'$, which was then used in Equation S8 to set CIs around the point estimate.

**Effective number of chromosomes and recombination**

Let $L_i$ be the length of the $i^{th}$ chromosome in bp, and let $L_T = \Sigma L_i$ be the total length of all chromosomes. Then $x_i = L_i/L_T$ is the relative (standardized) length of the $i^{th}$ chromosome, such that $\Sigma x_i=1$. It follows that the probability that two randomly drawn loci are both from chromosome $i$ is $x_i^2$, and the overall probability that both loci are drawn from the same chromosome is $\Sigma x_i^2$. The expected proportion of locus pairs that are on different chromosomes (unlinked) is therefore 1 - $\Sigma x_i^2$, which is easily calculated from the relative sizes of the chromosomes. In a way exactly analogous top the method for defining the effective number of alleles, we can define an effective number of chromosomes ($C_e$) as the number of equally-sized chromosomes that would produce the same expected fraction of unlinked (or linked) loci as the dataset in question. If there are $C$ equal-sized chromosomes, each one has relative size $x_i = 1/C$, so $\Sigma x_i^2 = C(1/C)^2 = 1/C$. The effective number of chromosomes is therefore easily calculated as

$$C_e = 1/\Sigma x_i^2. \quad \text{(S9)}$$

In our simulations, all chromosomes were the same size so the effective number of chromosomes was the same as the actual number. For other datasets, users can calculate $C_e$ and use that to compare with our simulated data.

**Example**: Hypothetical Species A has 20 chromosomes, 10 long and 10 short, with the long ones being 75 Mbp in length and the short ones 25. Then $L_T = \Sigma L_i = 10*25 + 10*75 = 1000$ Mbp. Each long chromosome therefore has $x_i = 75/1000 = 0.075$ and each short one has $x_i = 0.025$. The sum of squares is $\Sigma x_i^2 = 10*0.075^2 + 10*0.025^2 = 0.0625$, which leads to $C_e = 1/\Sigma x_i^2 = 16$. This means that for Species A, 6.25% of the pairwise comparisons should involve loci on the same chromosome, which is the same result you would get with 16 equal-sized chromosomes. This hypothetical example produces a $C_e/C$ ratio of 16/20 = 0.8. The $C_e/C$ ratio can vary widely among species; for example, based on data from NCBI for autosomes, we calculate $C_e/C$ = 0.84 in humans (Piovesan et al. 2019), 0.89 in dogs (https://www.ncbi.nlm.nih.gov/assembly/GCF_000002285.5#/st), and 0.42 in collared flycatchers (https://www.ncbi.nlm.nih.gov/assembly/GCF_000247815.1/#/st).

Veller et al. ( 2019) defined the metric $\bar{r}$ and showed that its expected value is given by a simple expression that has two components, one for independent assortment among chromosomes, and one for recombination within chromosomes. Making the substitution $\Sigma x_i^2 = 1/C_e$ based on the above results and converting to the notation used here, Equation 6 from Veller et al. (2019) can be expressed as:

$$E(\bar{r}) = \frac{1}{2}\sum_{i=1}^{C}\left(x_i^2 - \sum_{k=1}^{I_i+1} l_{i,k}^2\right) + \frac{1}{2}[1 - 1/C_e]. \quad \text{(S10)}$$

In this formulation, $I_i$ is the number of COs on bivalent chromosome $i$ (which divide the chromosome into $I_i$+1 segments), with the scaled length (relative to total genome length) of the $k^{th}$ segment being $l_{i,k}$. Focusing on the intra-chromosomal component,

$$\frac{1}{2}\sum_{i=1}^{C}\left(x_i^2 - \sum_{k=1}^{I_i+1} l_{i,k}^2\right) = \frac{1}{2}\left[\sum_{i=1}^{C} x_i^2 - \sum_{i=1}^{C}\sum_{k=1}^{I_i+1} l_{i,k}^2\right]$$
$$= \frac{1}{2}\left(\frac{1}{C_e}\right) - \frac{1}{2}\sum_{i=1}^{C}\sum_{k=1}^{I_i+1} l_{i,k}^2 ,$$

so Eq S10 can be further parsed as follows:

$$E(\bar{r}) = \frac{1}{2}\left(\frac{1}{C_e}\right) - \frac{1}{2}\sum_{i=1}^{C}\sum_{k=1}^{I_i+1} l_{i,k}^2 + \frac{1}{2}[1 - 1/C_e]$$
$$= 0.5 + \frac{1}{2C_e} - \frac{1}{2C_e} - \frac{1}{2}\sum_{i=1}^{C}\sum_{k=1}^{I_i+1} l_{i,k}^2$$
$$= 0.5 - \frac{1}{2}\sum_{i=1}^{C}\sum_{k=1}^{I_i+1} l_{i,k}^2 \quad , \quad \text{(S11)}$$

which is a slight variation of Eq 3 in Veller et al. 2019. This equation makes no assumptions about the relative chromosome lengths, $x_i$. In our model, all $x_i = 1/C$. If we let the length of each chromosome be $L_i$ = 1 unit, then total genome length is $L_T = C$ and all the $l_{i,k}$ terms can be expressed as $l_{i,k} = q_{i,k}/C$, where $q_{i,k}$ is the actual (unscaled) length of segment $k$ within chromosome $i$. Within each chromosome, $\Sigma q_{i,k} = 1$. In our model, then,

$$E(\bar{r}) = 0.5 - \frac{1}{2}\frac{1}{C^2}\sum_{i=1}^{C}\sum_{k=1}^{l_i+1} q_{i,k}^2. \quad \text{(S12)}$$

The right side of this equation has $C$ terms, each of which is the sum of $q_{i,k}^2$ across the number of segments within one chromosome. If we let $\overline{Q^2}$ be the mean of these terms, then Eq S12 can be written as

$$E(\bar{r}) = 0.5 - \frac{1}{2C^2} C * \overline{Q^2} = 0.5 - \frac{\overline{Q^2}}{2C} \; . \quad \text{(S13)}$$

Note that in our model, $\overline{Q^2}$ is independent of the number of chromosomes, so we can evaluate $\overline{Q^2}$ using a simple model with 1 chromosome. This is easy to do by simulation, using code that is provided here (https://github.com/nwfsc-cb/pseudorep). Our simulations assumed that the number of COs per chromosome was Poisson distributed with a mean of 0.5, and that COs were randomly spaced along the chromosome (hence no interference). Under these conditions, we find that $\overline{Q^2}$ = 0.852, which leads to

$$E(\bar{r}) = 0.5 - \frac{\overline{Q^2}}{2C} = 0.5 - \frac{0.852}{2C} \; . \quad \text{(S14)}$$

Equation S14 provides a means whereby researchers interested in a particular species (Species B) can use our framework to account for the intra-chromosomal component of $\bar{r}$, in addition to the effects of variable chromosome size. The steps are as follows:

1. For Species B, compute the effective number of chromosomes, $C_e$, based on variation in chromosome length.
2. Use the value of $C_e$ for $C$ in Eq S14 to calculate $E(\bar{r})$ under our model. The result is the total expected amount of genetic shuffling that would occur under our model, due both to unequal chromosome size and recombination within chromosomes.
3. For Species B, estimate the actual $E(\bar{r})$, using Equations 3A, S10, or S11. Alternatively, conduct a sensitivity analysis using various adjustments to how recombination is treated in our model.
4. Find the number of equal-sized chromosomes ($C_{eAdj}$) which, when inserted into Eq S14, produces the correct $E(\bar{r})$ for Species B. This can be done using Eq S15, below.
5. Use our model-fitting results with effective number of chromosomes = $C_{eAdj}$ to estimate effective df for user-specified values of $N_e$, $L$, and $S$.

To illustrate, assume that the focal species has $C_e$ = 16 effective chromosomes, based only on variation in chromosome size. Inserting $C$ = 16 into Equation S14 produces $E(\bar{r})$ = 0.5 – 0.852/32 = 0.4734. Of this total, the inter-chromosomal effect (second term in Eq S10) accounts for 0.5(1-1/16) = 0.4688 (99% of the total genetic shuffling) and recombination within chromosomes (0.0046) accounts for the remaining 1%. Assume now that in Species B, the recombination rate is twice what we modeled (mean = 1 per chromosome rather than 0.5). In our model this would cut within-chromosome segments into smaller fragments, with the result that $\overline{Q^2}$ = 0.735 and $E(\bar{r})$ = 0.5 – 0.735/32 = 0.4770 (this result can be replicated by changing the mean number of COs in our code mentioned above).

For Step 3 above, we want to find the number of equal-sized chromosomes which, in our modeling system, would produce the actual amount of genetic shuffling in Species B. The general relationship is:

$$E(\bar{r})_{SpeciesB} = 0.5 - \frac{0.852}{2C_{eAdj}} \quad ,$$

and rearranging and solving for $C$ gives

$$C_{eAdj} = \frac{0.852/2}{0.5 - E(\bar{r})_{SpeciesB}}\ . \qquad \text{(S15)}$$

Substituting the estimated value for $E(\bar{r})_{SpeciesB}$ in Eq S15 produces $C_{eAdj}$ = (0.852/2)/(0.5-0.477) = 18.5. Thus, under our model a two-fold increase in the recombination rate would increase the intra-chromosomal effect from 0.0046 to 0.0082, would increase the overall amount of genetic shuffling from 0.4734 to 0.4770, and would increase the adjusted effective number of chromosomes from 16 to 18.5, an increase of 16%.

We can evaluate the consequences of this increased amount of recombination for pseudoreplication by using our model to estimate the change in effective df. For illustration, assume that $S$=50 individuals of Species B were sampled from each of two isolated populations with $N_e$=200, and that mean $F_{ST}$ was calculated using $L$ = 10000 SNPs. Using our model-fitting code (available at https://github.com/nwfsc-cb/pseudorep), $L$' for $F_{ST}$ is 5035 for 16 chromosomes and 5416 for 18.5 chromosomes, an increase of 8% after accounting for a doubling of the overall recombination rate. For LD based on mean $r^2$ for individual samples, the comparable values are $n$' = 14031 for $C$ = 16 and 14583 for $C$ = 18.5, an increase of 4%. As expected, we see a proportionally smaller effect on pseudoreplication for LD, because overlapping pairs of loci (which are not directly affected by physical linkage or recombination) are an important contributor to lack of independence of pairwise $r^2$ values.

**Juvenile sampling**

Our simulations modeled hypergeometric sampling of $S$ individuals drawn without replacement from the $N_e$ = $N$ total in the population. If $P$ is the frequency of an allele in the overall population, then the variance of allele frequency in the sample is given by

$$Var(\hat{P}) = \frac{P(1-P)}{2S}(N-S)/N = P(1-P)\frac{N-S}{2NS}\ . \qquad \text{(S16)}$$

If instead the parents are allowed to reproduce and juvenile offspring are sampled, this variance becomes (Nei and Tajima 1981):

$$Var(\hat{P}) = P(1-P)\left[1-\left(1-\frac{1}{2N_e}\right)\left(1-\frac{1}{2S}\right)\right].$$

Ignoring the term 1/(4$SN_e$) because it will be small, this is closely approximated by

$$Var(\hat{P}) \approx P(1-P)\left[\left(\frac{1}{2N_e}\right)+\left(\frac{1}{2S}\right)\right] = P(1-P)\frac{N+S}{2NS}. \qquad \text{(S17)}$$

It is easy to see that the two sampling variances converge as $S$ becomes small relative to $N_e$. Therefore, a robust estimate of precision for juvenile sampling can be obtained by using the predicted effective df from this study for small $S$.

**References** (showing only those not cited in main text)


Bolker B and R Development Core Team (2020). bbmle: Tools for General Maximum Likelihood Estimation. R package version 1.0.23.1. https://CRAN.R-project.org/package=bbmle.

Booker, T.R., Yeaman, S. and Whitlock, M., 2020. Variation in recombination rate affects detection of FST outliers under neutrality. *bioRxiv*.

Do, C., R.S. Waples, D. Peel, G.M. Macbeth, B.J. Tillet, and J.R. Ovenden. 2014. NeEstimator V2: re-implementation of software for the estimation of contemporary effective population size (Ne) from genetic data. Molecular Ecology Resources 14:209-214 (DOI: 10.1111/1755-0998.12157).

Ewens WJ, Feldman MW. 1976. The theoretical assessment of selective neutrality. Pp. 303-338 in: Population Genetics and Ecology. S Karlin and E Nevo, eds. Academic Press, New York.

Gao XY, Stamier J, Martin ER. 2008. A multiple testing correction method for genetic association studies using correlated single nucleotide polymorphisms. *Genetic Epidem*. 32:361-369.

Kelleher, J., Thornton, K.R., Ashander, J. and Ralph, P.L., 2018. Efficient pedigree recording for fast population genetics simulation. *PLoS computational biology*, *14*(11), p.e1006581.

Kosambi, D.D. 1944. The estimation of map distance from recombination values, *Annals of Eugenics*, 12: 172–175.

Lotterhos, K.E., 2019. The effect of neutral recombination variation on genome scans for selection. *G3: Genes, Genomes, Genetics*, *9*(6), pp.1851-1867.

Nei M, Maruyama T. 1975. Lewontin-Krakauer test for neutral genes. Genetics 80:395.

Piovesan, A., Pelleri, M.C., Antonaros, F., Strippoli, P., Caracausi, M. and Vitale, L., 2019. On the length, weight and GC content of the human genome. *BMC research notes*, *12*(1), pp.1-7.

Pollak, E., 1983 A new method for estimating the effective population size from allele frequency changes. Genetics 104:531-548.

Pya N (2020). scam: Shape Constrained Additive Models. R package version 1.2-6. https://CRAN.R-project.org/package=scam

Reich D, Thangaraj K, Patterson N, Price AL, Singh L. 2009. Reconstructing Indian population history. Nature 461(7263):489–494.

Robertson A. 1975. Gene frequency distribution as a test of selective neutrality. Genetics 81:775-785.

Sved JA, Feldman MW (1973). Correlation and probability methods for one and two loci. Theor Popul Biol 4: 129–132.

Weir BS, Hill WG (1980). Effect of mating structure on variation in linkage disequilibrium. Genetics 95: 477–488.

Wood, S.N. (2004) Stable and efficient multiple smoothing parameter estimation for generalized additive models. Journal of the American Statistical Association. 99:673-686.

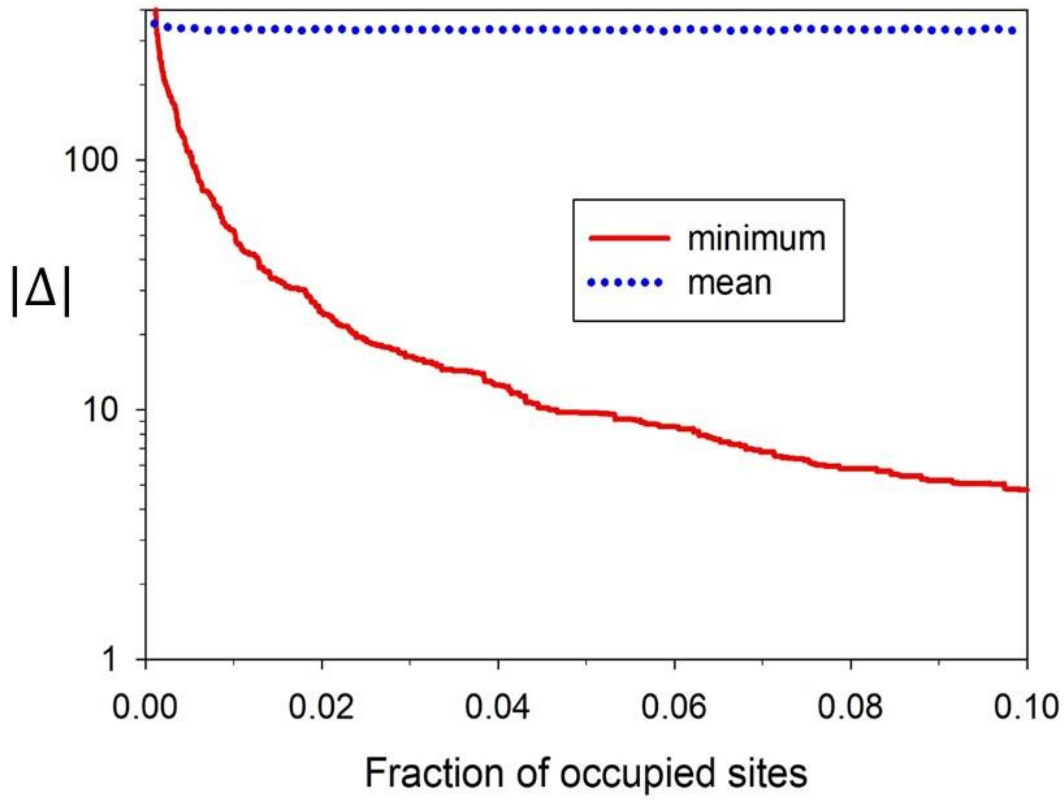


Figure S1. Absolute value of the distance (Δ) between a new gene locus being added to a chromosome and existing loci, as a function of the fraction of sites that are already occupied. Possible sites for loci are evenly spaced across the chromosome one unit apart. Mean |Δ| between the new locus and all existing pairs does not change with the number of loci, but minimum |Δ| declines. These results are for a simulation with 10,000 loci randomly sprinkled on a chromosome of length 100,000 units, but the pattern does not depend on specific numbers of loci and chromosome length—only their ratio.

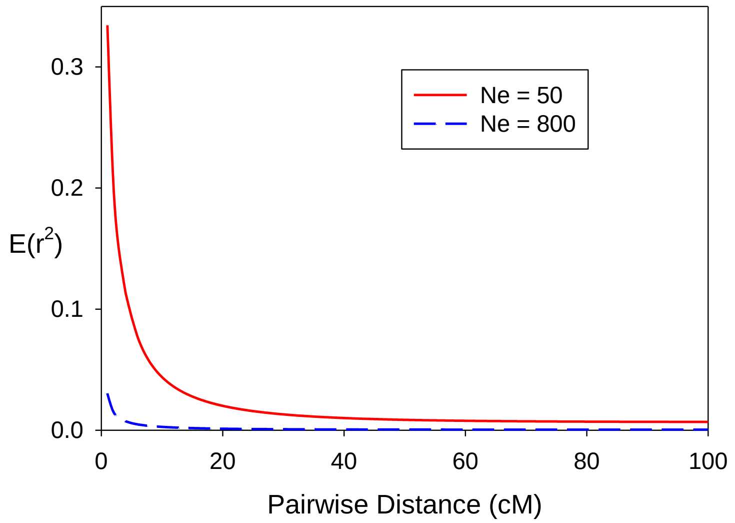


Figure S2. Rate of decay of LD as a function of $N_e$ and the distance in centimorgans between a pair of SNPs. We assumed a Kosambi function (Kosambi 1944) relating distance to recombination fraction ($c$) and a hybrid Sved/Feldman – Weir/Hill relationship between E($r^2$), $c$, and $N_e$.

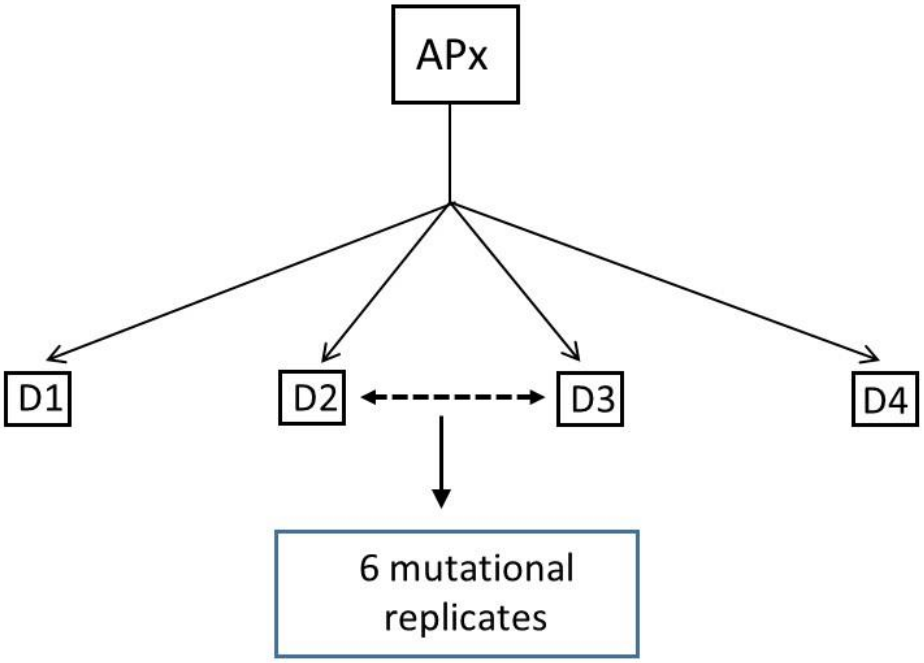


Figure S3. Experimental design for simulations involving $F_{ST}$. For each evolutionary scenario (combination of $N_e$ and genome size), four ancestral populations (AP1–AP4; only one of which, APx, is shown here) were simulated to ensure coalescence ($10N_e$ generations). Next, each ancestral population split into four daughter populations (D1-D4), which then evolved independently under isolation for another $t = 0.2N_e$ generations. In the final generation, six pairwise comparisons of the four daughter populations are possible (only that between D2 and D3 is shown here), with each pair representing a different two-population pedigree. Six replicate sets of genotypes were then generated by adding mutations to gene trees determined by each population pedigree, and replicate samples were then taken to compute $F_{ST}$ and $n$'.

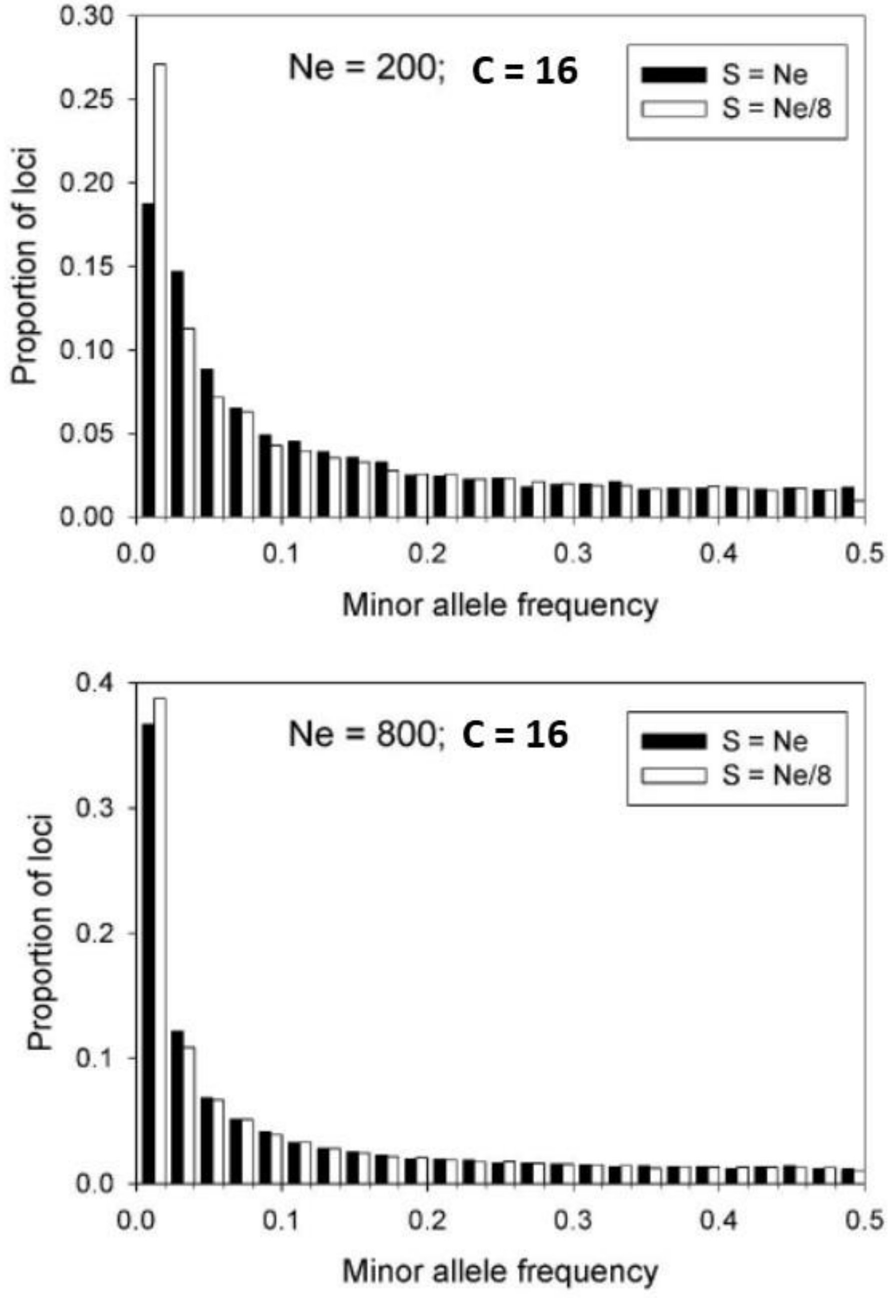


Figure S4. Distribution of minor allele frequency at the final generation for representative scenarios from the LD simulations. Results are shown for 50K randomly-selected SNPs for samples of size $S = N_e$ (sampling the entire population) and $S = N_e/8$.

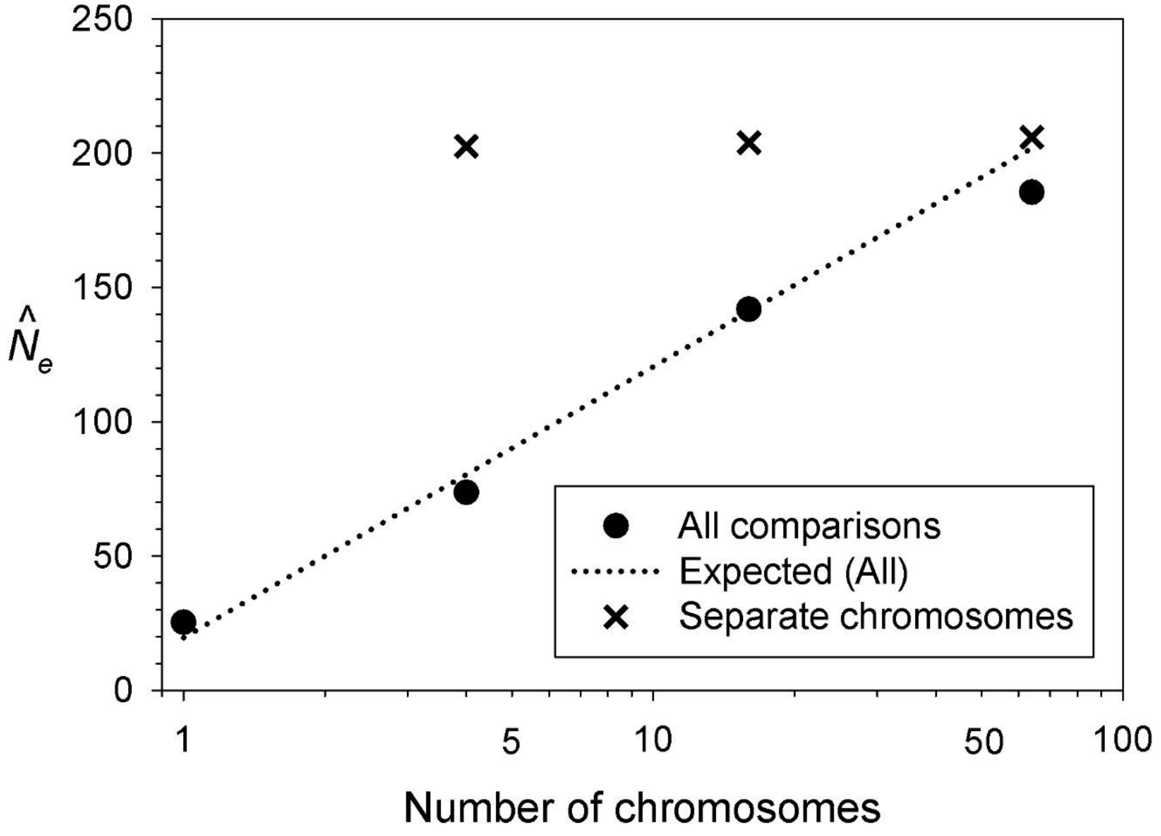


Figure S5. Estimates of $N_e$ from the LDNe method (Waples and Do 2008), based on mean $r^2$ values computed over all pairs of loci (filled circles) or only pairs on different chromosomes (Xs). Shown are results of simulations with true $N_e = 200$, $S = 200$, and up to 75,000 loci randomly distributed on 1, 4, 16, or 64 equal-sized chromosomes. The dotted line shows $E(\widehat{N}_e)$ based on the bias adjustment proposed by Waples et al. (2016).

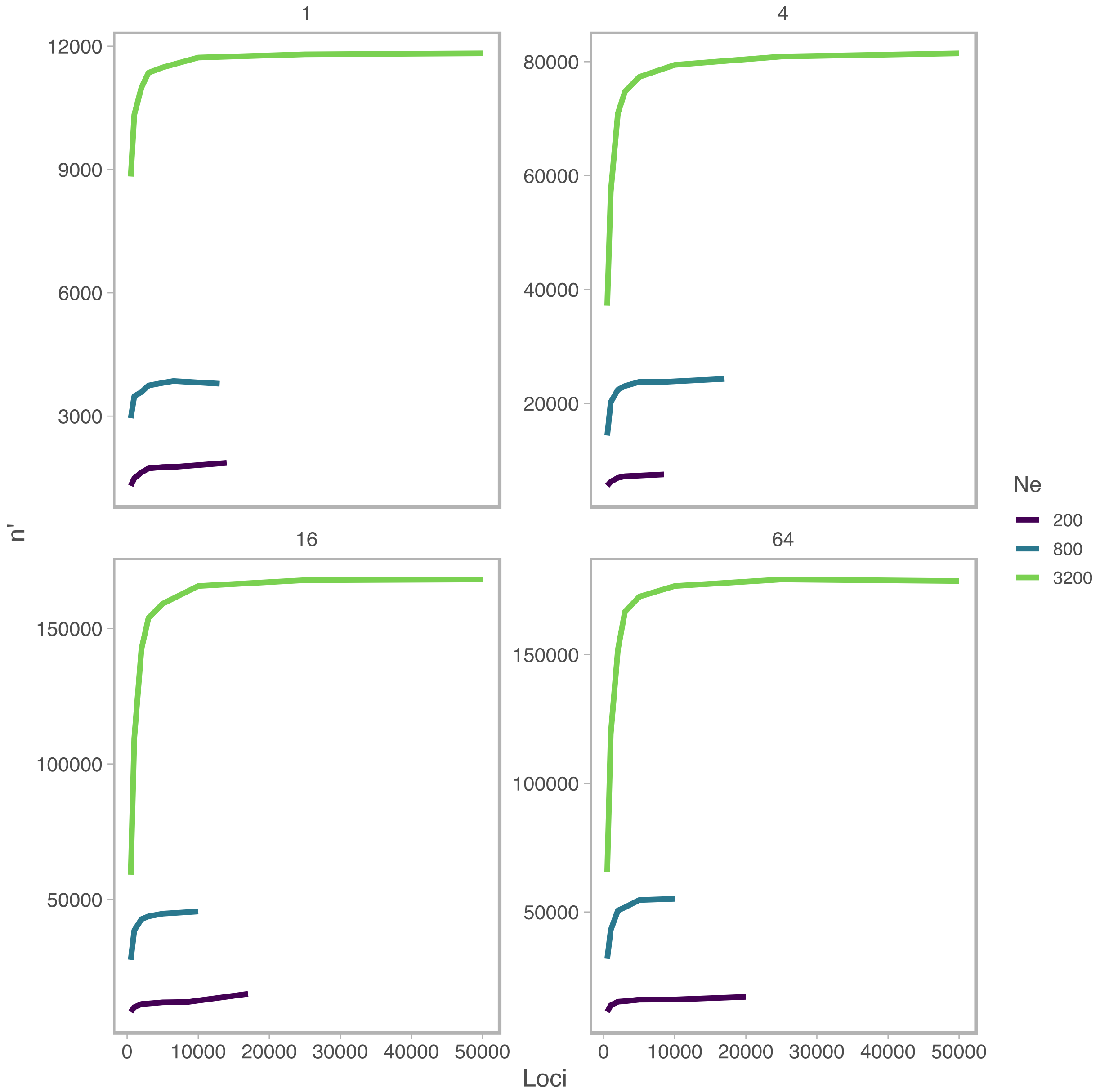


Figure S6. Simulated data for LD analyses. These figures show the relationship between the effective df ($n'$) and the number of loci ($L$) and $N_e$, and the panels are faceted by chromosome number, $C$ = [1,4,16,64] (allowing a different y-axis for each).

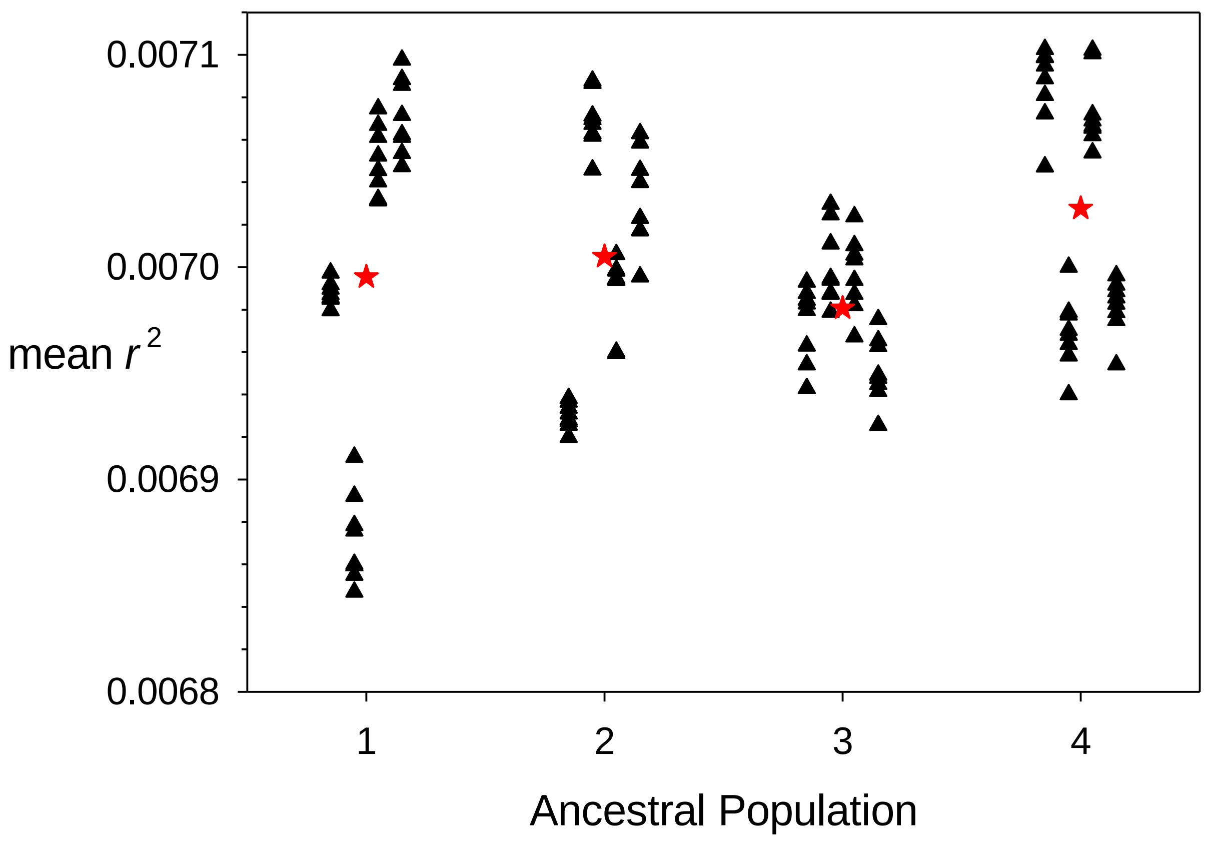


Figure S7. Distribution of mean $r^2$ calculated across all pairs of 5000 diallelic loci, for 128 replicate populations simulated with $N_e = 200$ and 64 chromosomes. For each of 4 ancestral populations, 4 isolated daughter populations were simulated, 8 mutational replicates were generated for each daughter population (see Figure 1), and mean $r^2$ was calculated across all 200 individuals in the final generations. Each vertical set of eight black triangles shows results for different mutational replicates for a single daughter population, which are jittered to avoid overlap; red stars are means across all datasets descended from a single ancestral population.

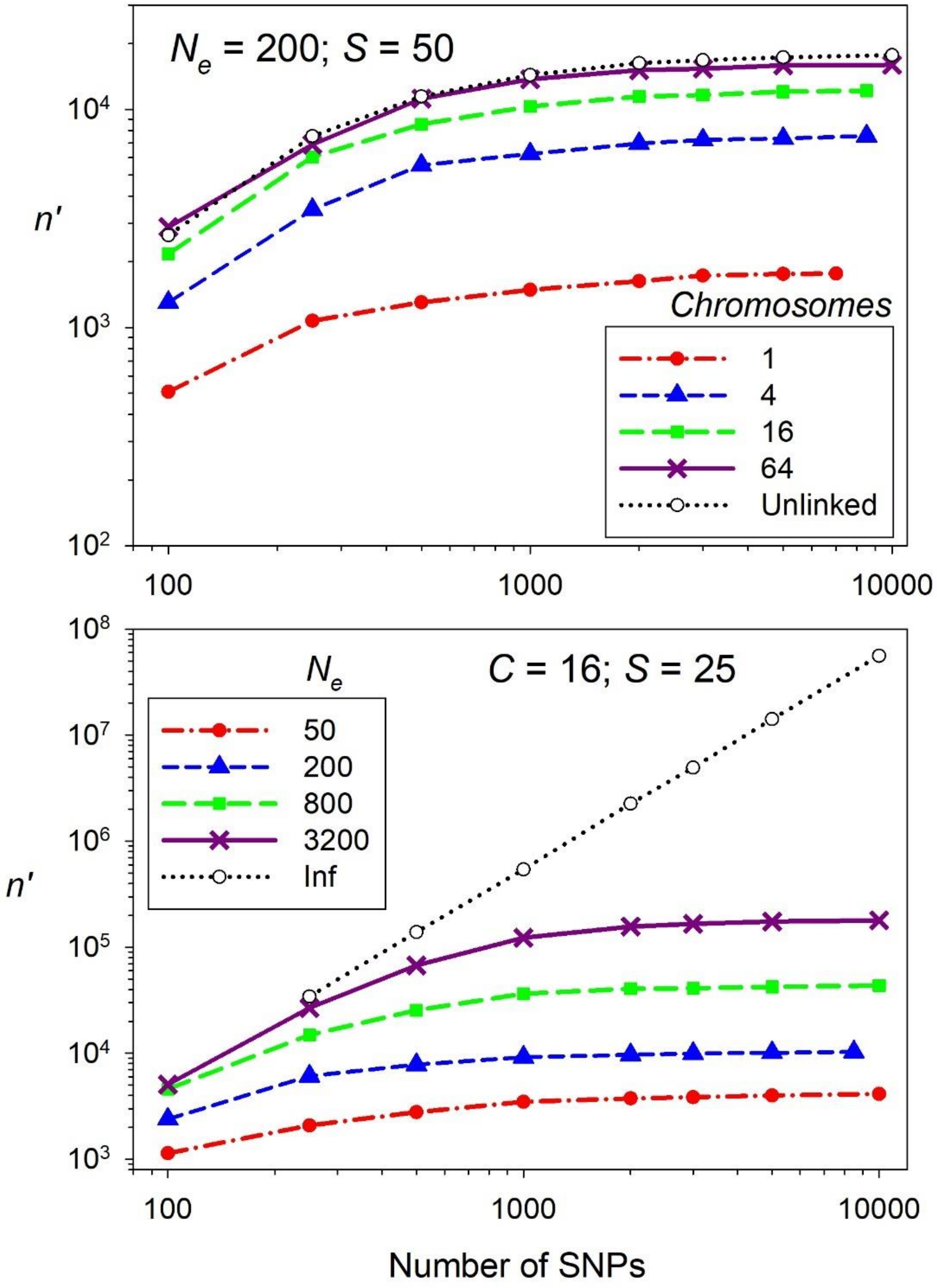


Figure S8. Effective degrees of freedom ($n$') for mean $r^2$ as a function of the number of diallelic (SNP) loci, $L$. Top: Effect of number of chromosomes ($C$), with $N_e$ = 200 and $S$ = 50. Bottom: effect of $N_e$, with $C$ = 16 and $S$ = 25. Mean $r^2$ was calculated across all $n$ pairs of loci. Figure 2 (main text) shows these same results except the Y axis is plotted as the effective number of loci ($L$').

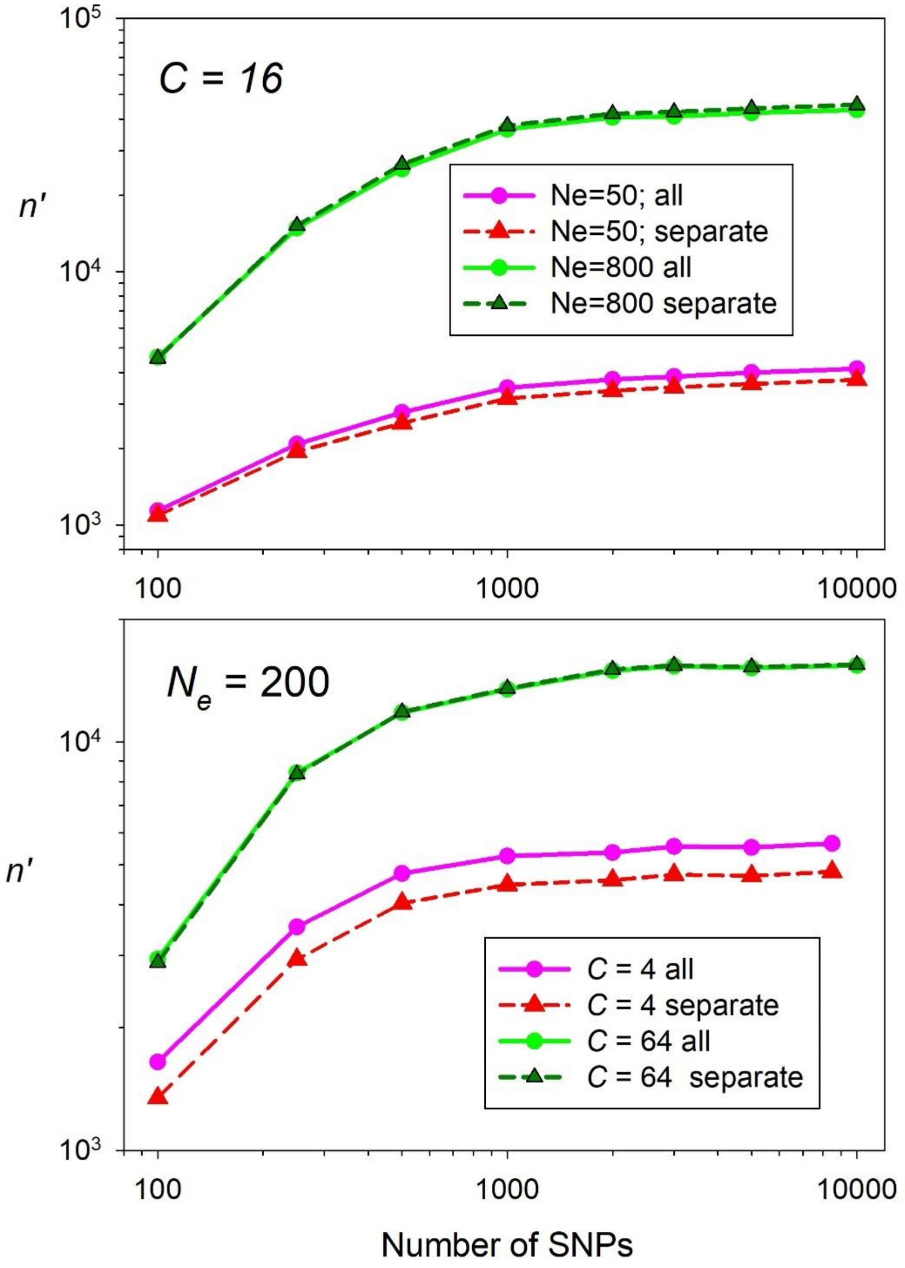


Figure S9. Effective degrees of freedom ($n$') for mean $r^2$ as a function of number of diallelic (SNP) loci and whether mean $r^2$ is calculated across all pairs of loci or only those on separate chromosomes. Top: $N_e$ = [50,800] with 16 chromosomes ($C$). Bottom: $C$ = [4,64] with $N_e$ = 200. Sample size of individuals was $S$ = 25.

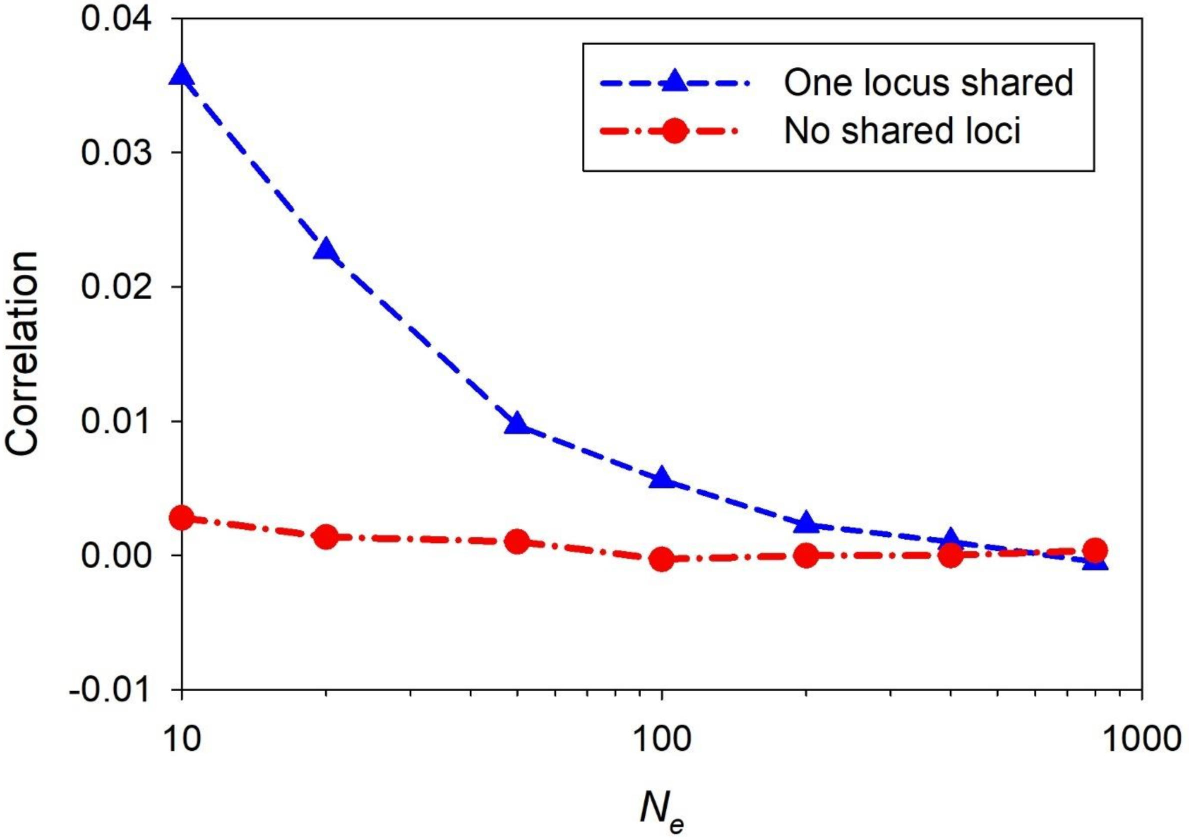


Figure S10. Mean correlations between pairwise $r^2$ values that do or do not share one locus. Results are for simulations where $r^2$ was computed across $N_e$ = 10-800 individuals for all 45 pairs of 10 unlinked, diallelic (SNP) loci.

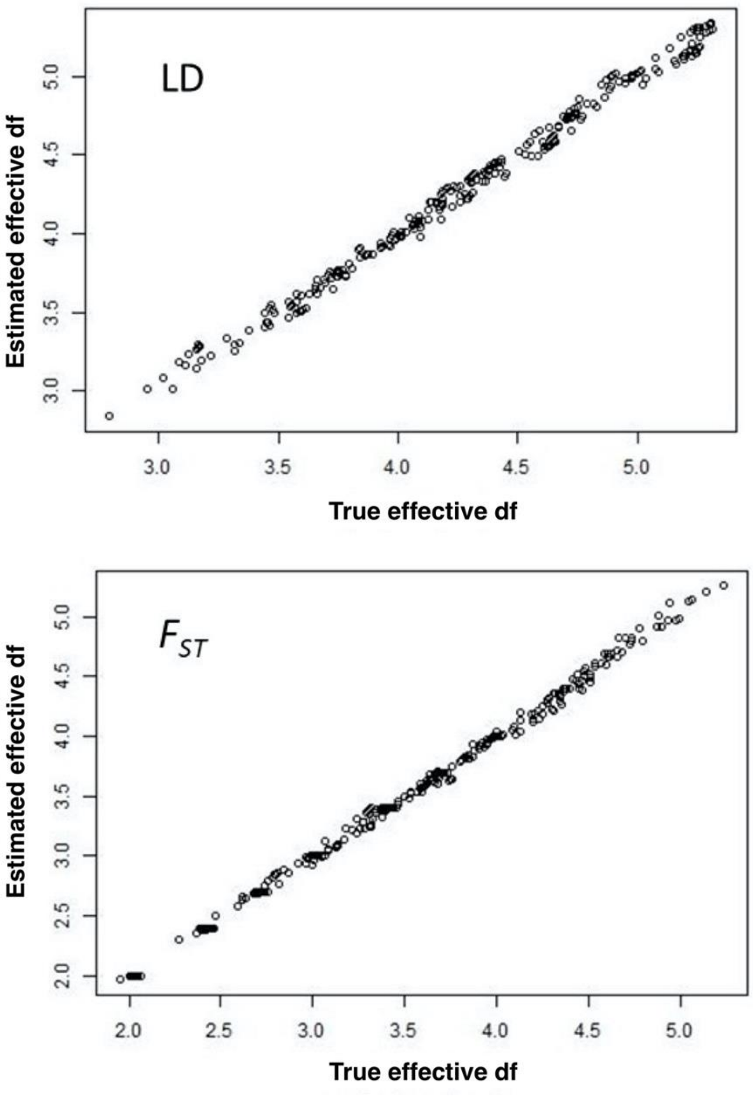


Figure S11. Diagnostic plots, showing predicted versus observed, with log(effective df) as the response. For LD, effective df = *n'* = effective number of pairs of loci; for $F_{ST}$, effective df = *L'* = effective number of loci. Top: LD analyses; correlation between the observed and predicted is 0.997. Bottom: $F_{ST}$: correlation between the observed and predicted is 0.998.

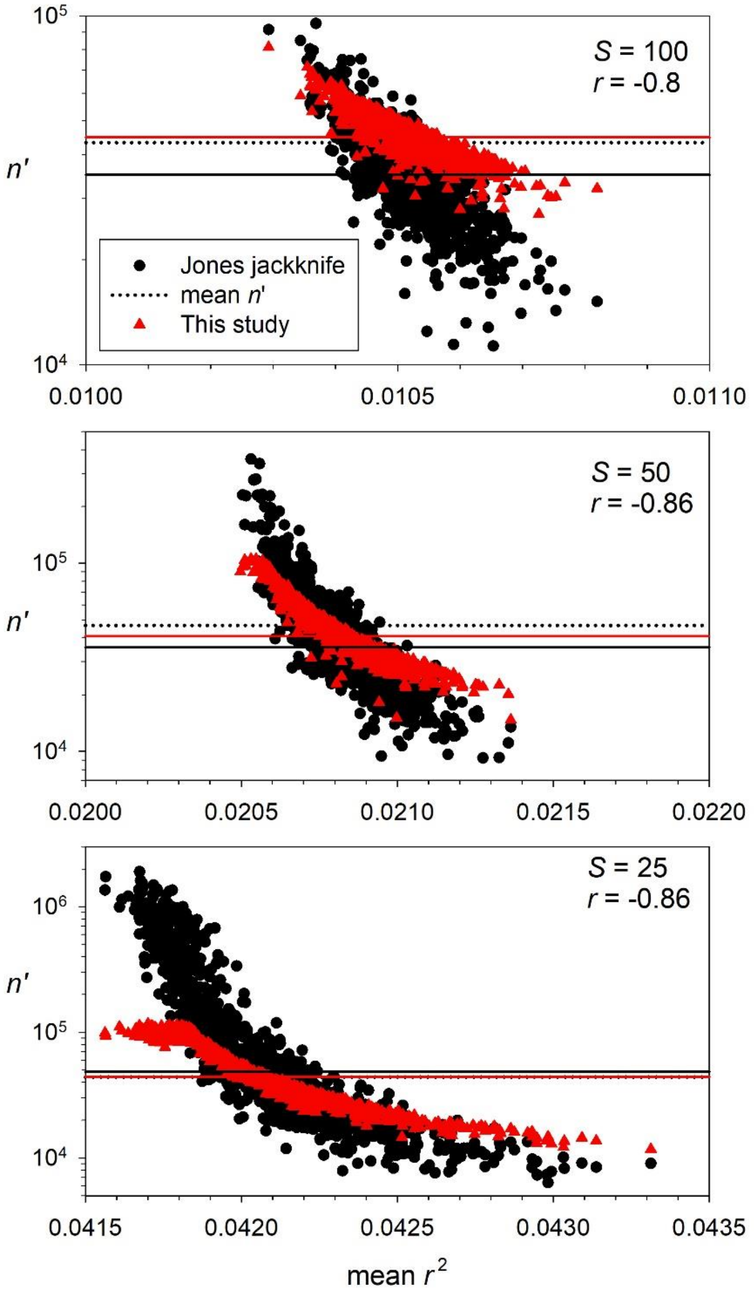


Figure S12. Scatterplots showing correlation between estimated $n$' and mean $r^2$ for LD analyses. Black circles are for the Jones et al. (2016) jackknife method and red triangles used methods described in this paper. Each datapoint is from one of 1024 replicate samples, generated using $N_e = 800$, $C = 16$, $S = 25$-100, and pairwise comparisons of 5000 diallelic (SNP) loci that are on different chromosomes. Horizontal lines: black dotted = $n$' calculated in this study using a separate set of simulated data; black solid: median of the black symbols; red solid: median of red symbols.

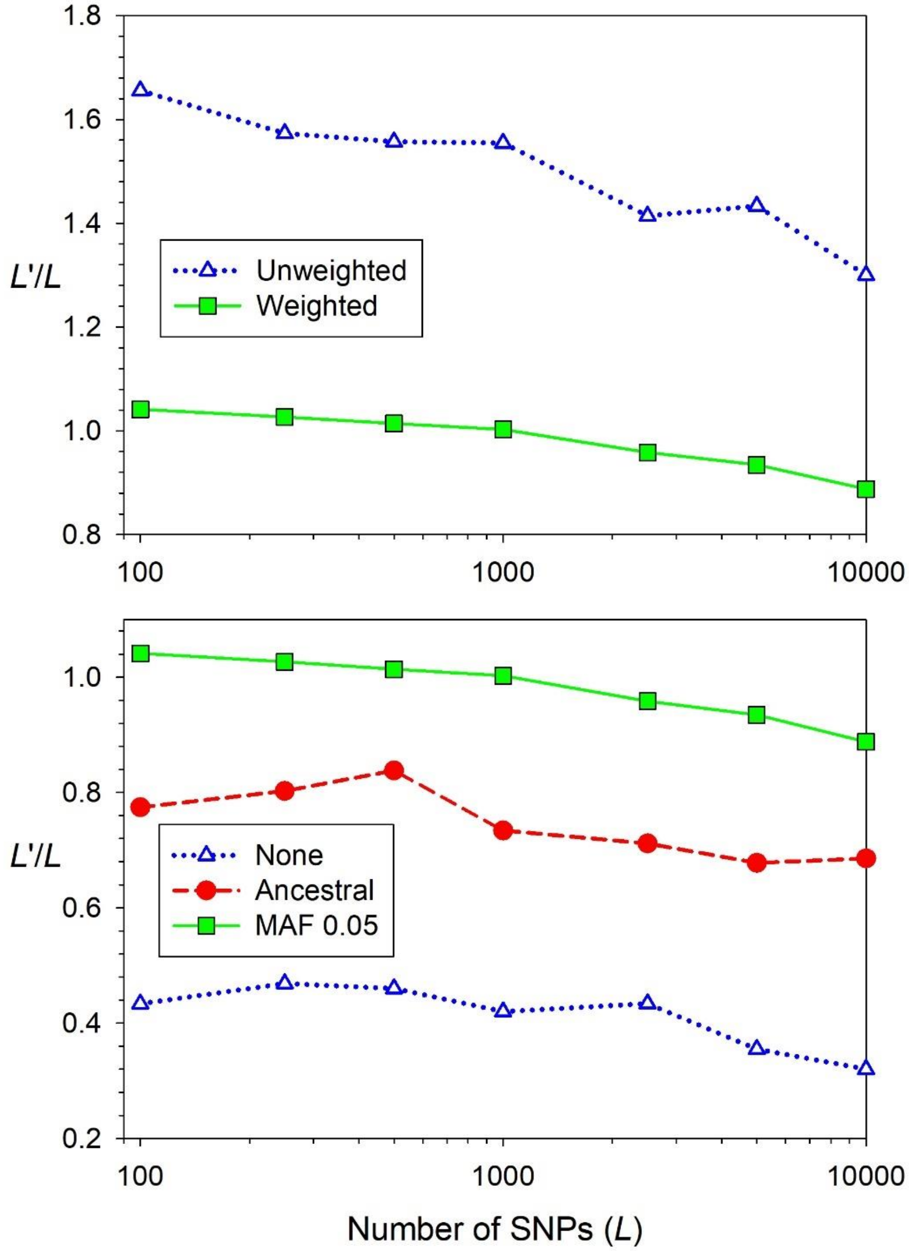


Figure S13. Sensitivity testing for various ways of computing $F_{ST}$. The $Y$ axis shows the ratio of effective degrees of freedom ($L$') to the number of diallelic (SNP) loci ($L$). Top panel: results for unweighted and weighted versions of $\hat{F}_{ST}^{Nei}$, using a MAF ≥ 0.05. Bottom panel: results for three ascertainment schemes, all using weighted $\hat{F}_{ST}^{Nei}$ ("none" = no ascertainment, all variable loci used; "ancestral" = ascertained in the ancestral population; "MAF 0.05" = used only loci with combined MAF ≥ 0.05 in samples analyzed). All results are for simulations with $N_e$ = 200, $C$ = 64 chromosomes, and $S$ = 50.

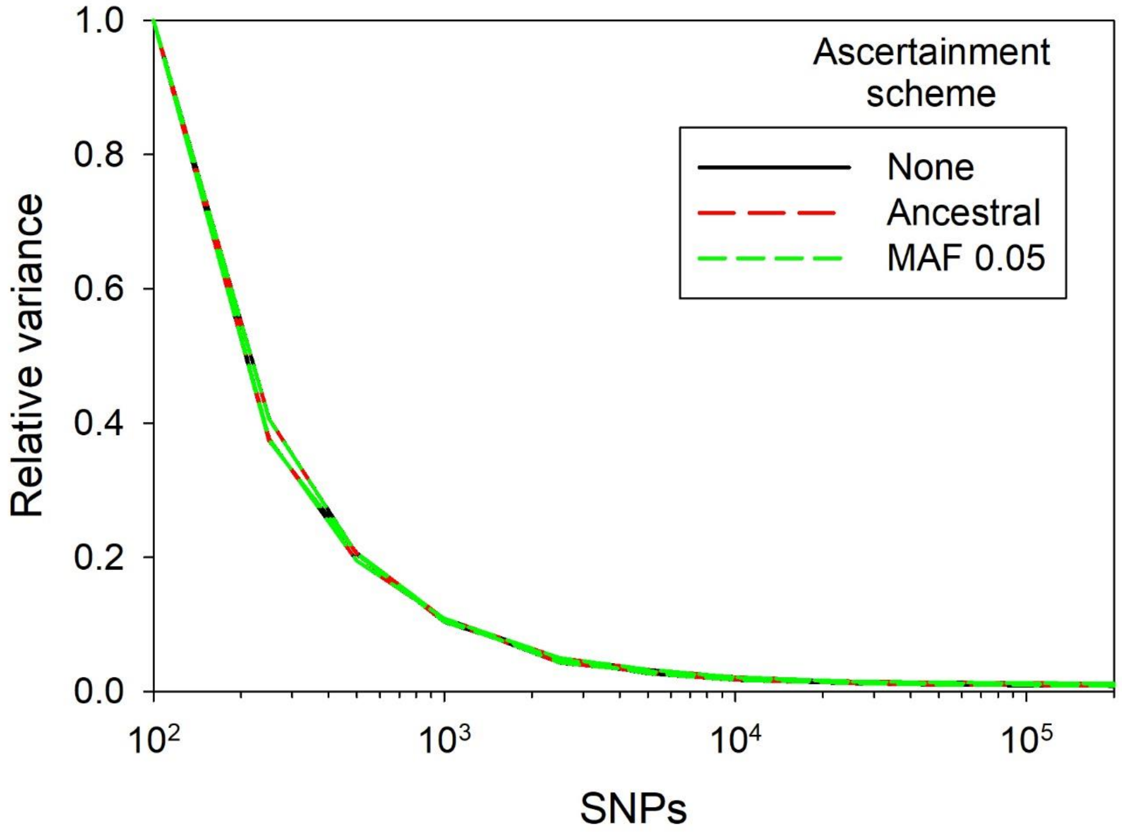


Figure S14. Rate of decline in var($\hat{F}_{ST}^{Nei}$) with increasing numbers of loci under 3 ascertainment schemes ("none" = no ascertainment, all variable loci used; "ancestral" = ascertained in the ancestral population; "MAF 0.05" = used only loci with combined MAF ≥ 0.05 in samples analyzed). Although the method of acertainment affects mean $\hat{F}_{ST}$ (see Figure S13), it does not affect the variance structure and hence calculation of *L*'. Results are for simulations with $N_e$ = 200, *C* = 16, and *S* = 50.

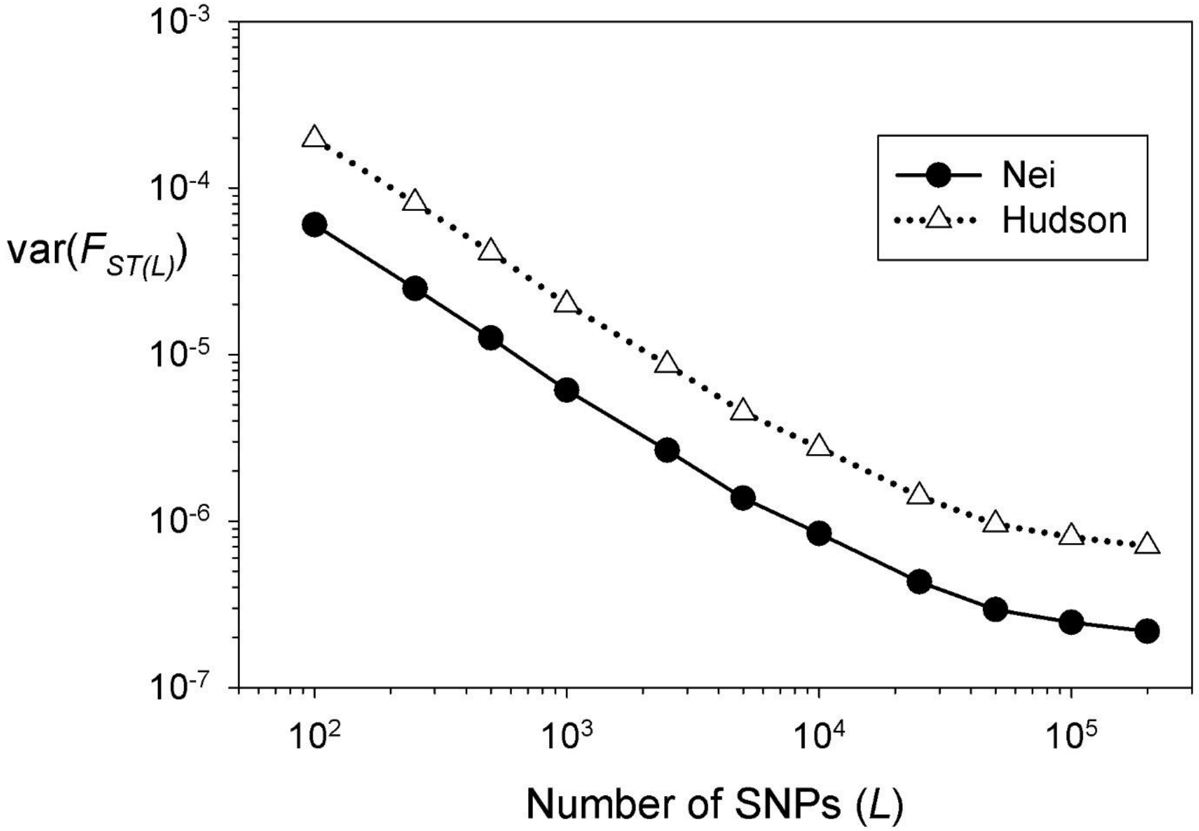


Figure S15. Rate of decline in the variance of multilocus $\hat{F}_{ST(L)}$ as more diallelic loci (SNPs, $L$) were used in the analysis. Results are for simulations with $N_e$ = 800, $C$ = 16, and $S$ = 50 and are shown for the estimators of Nei ($\hat{F}_{ST}^{Nei}$) and Hudson ($\hat{F}_{ST}^{Hudson}$). See Figure 6 (main text) for comparable results for a different scenario.

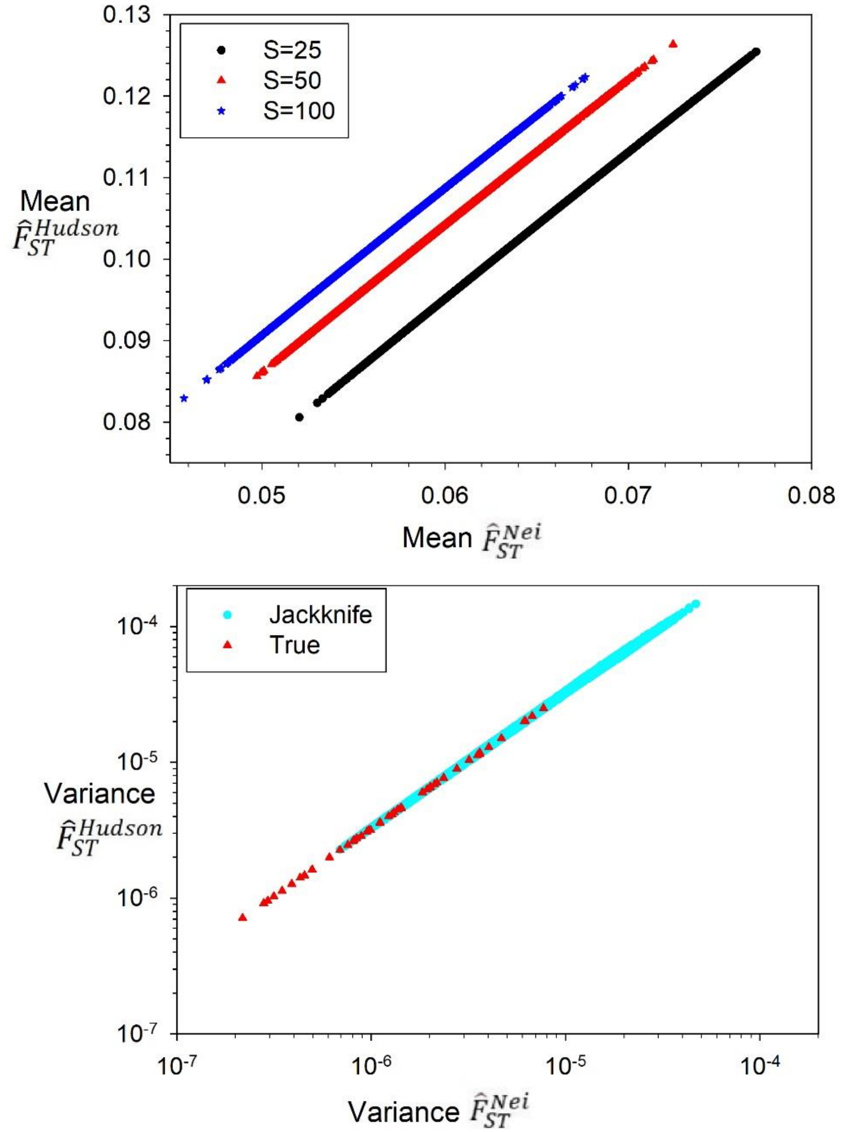


Figure S16. The Nei and Hudson estimators are almost perfectly correlated ($r^2 > 0.999$ for all comparisons shown here). Top: mean values of the two estimators across >41K samples used for jackknife evaluations. For a given sample size, the two estimators are related by a common slope ($\hat{F}_{ST}^{Hudson} = 1.801\hat{F}_{ST}^{Nei}$) and a sample-size specific intercept (α = [-0.0129, -0.003842, -0.0006804] for $S$ = [25, 50, 100], respectively). Alternatively, the relationship can be expressed as $\hat{F}_{ST}^{Nei} = 0.555\hat{F}_{ST}^{Hudson}$ + [0.0072, 0.0021, 0.00037]. Bottom: Variance of $\hat{F}_{ST}^{Hudson}$ as a function of the variance of $\hat{F}_{ST}^{Nei}$. A single slope applies across all sample sizes and is nearly identical whether it compares 5 Mb block jackknife estimates of the variance ($\beta = 3.231$; filled cyan circles) or 'true' variances computed across many replicate samples derived from the same population pedigree ($\beta = 3.258$; red triangles). All data shown here are for simulations with $N_e$ = 200 or 800; 4 or 16 chromosomes; $L$ = 5K-50K loci; and sample sizes of $S$ = 25, 50, or 100 individuals. Slopes and intercepts relating the Hudson and Nei estimators are functions of mean $\hat{F}_{ST}^{Hudson}$ and mean $\hat{F}_{ST}^{Nei}$ (data not shown).

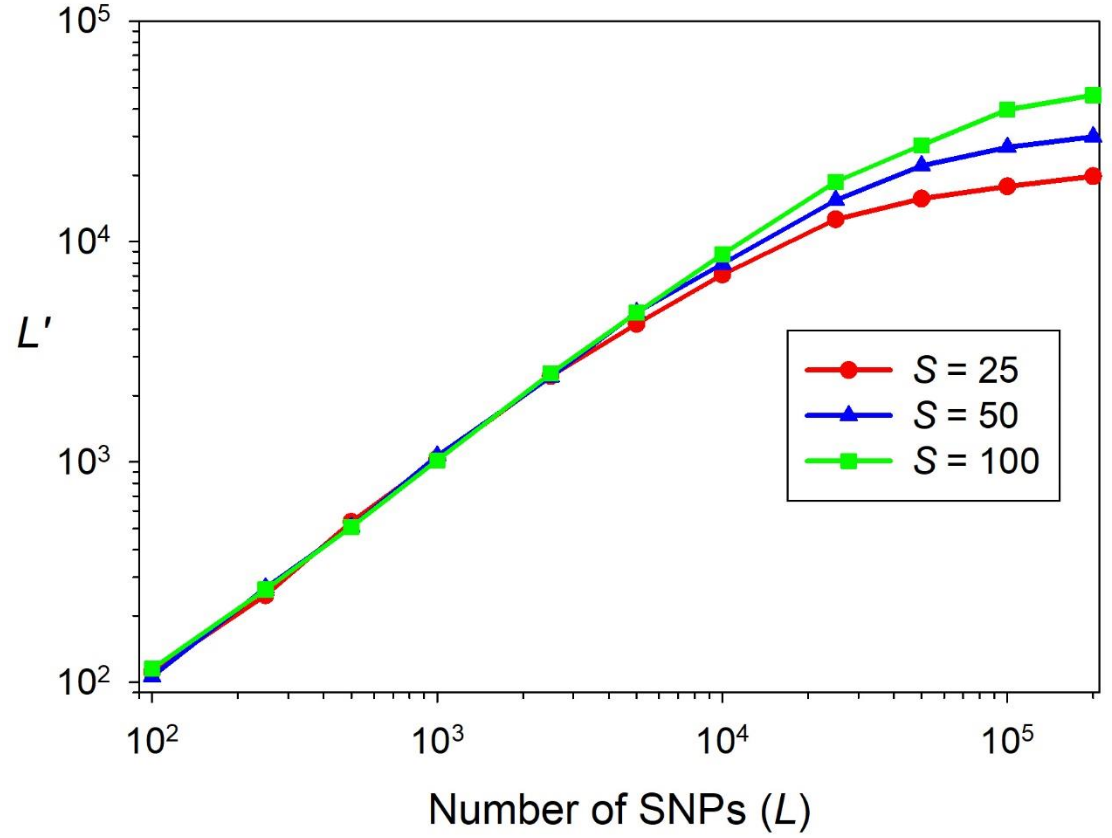


Figure S17. Effective degrees of freedom ($L$') for mean $F_{ST}$ as a function of the sample size of individuals ($S$ = 25-100) and the number of diallelic (SNP) loci, $L$. Results are for $N_e$ = 800, $C$ = 16.

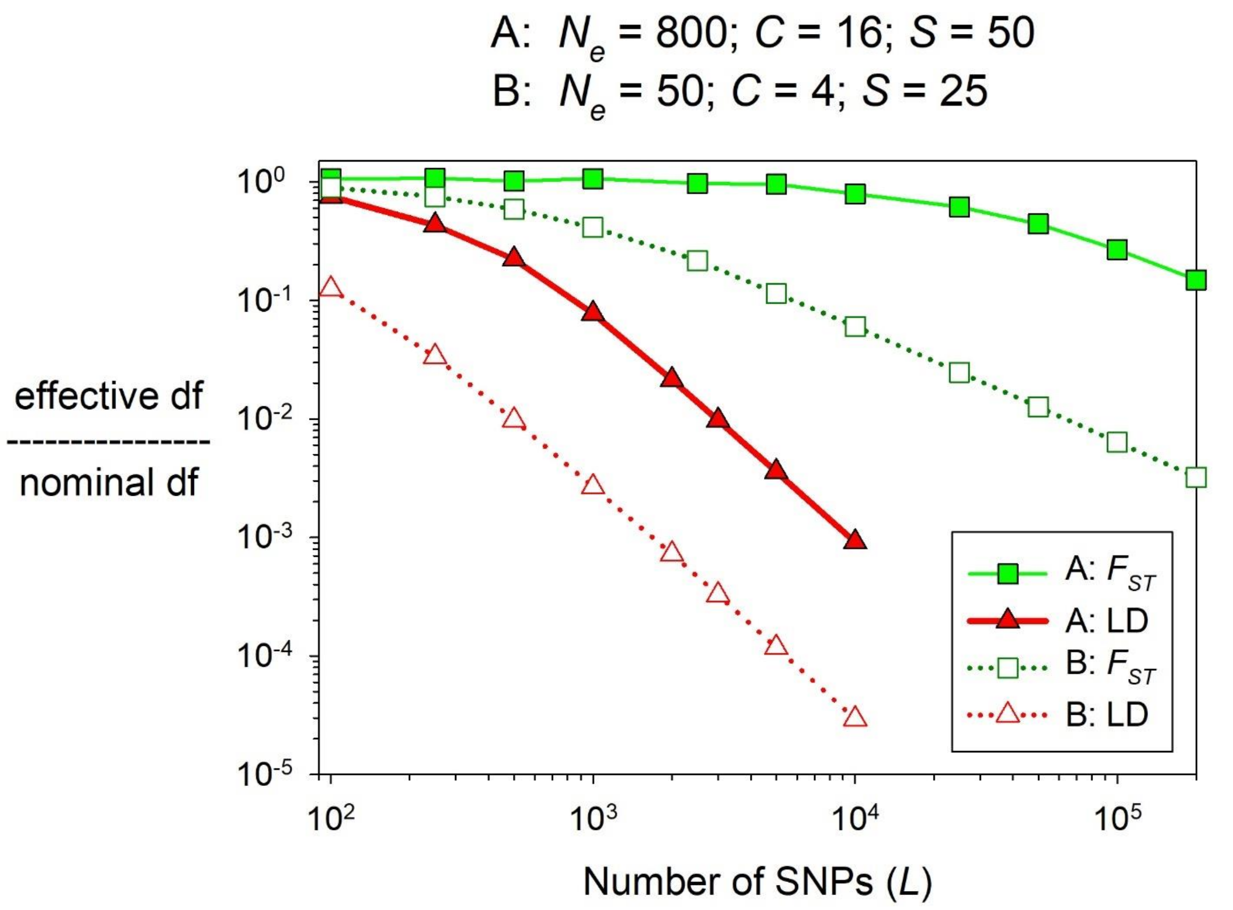


Figure S18. Comparison of the ratio of effective degrees of freedom (df) to nominal degrees of freedom for $F_{ST}$ and LD, for two different scenarios (A and B, with parameters as indicated in the figure). For $F_{ST}$, nominal df = $L$ = the number of diallelic (SNP) loci; for LD, nominal df = $n$ = $L(L-1)/2$.

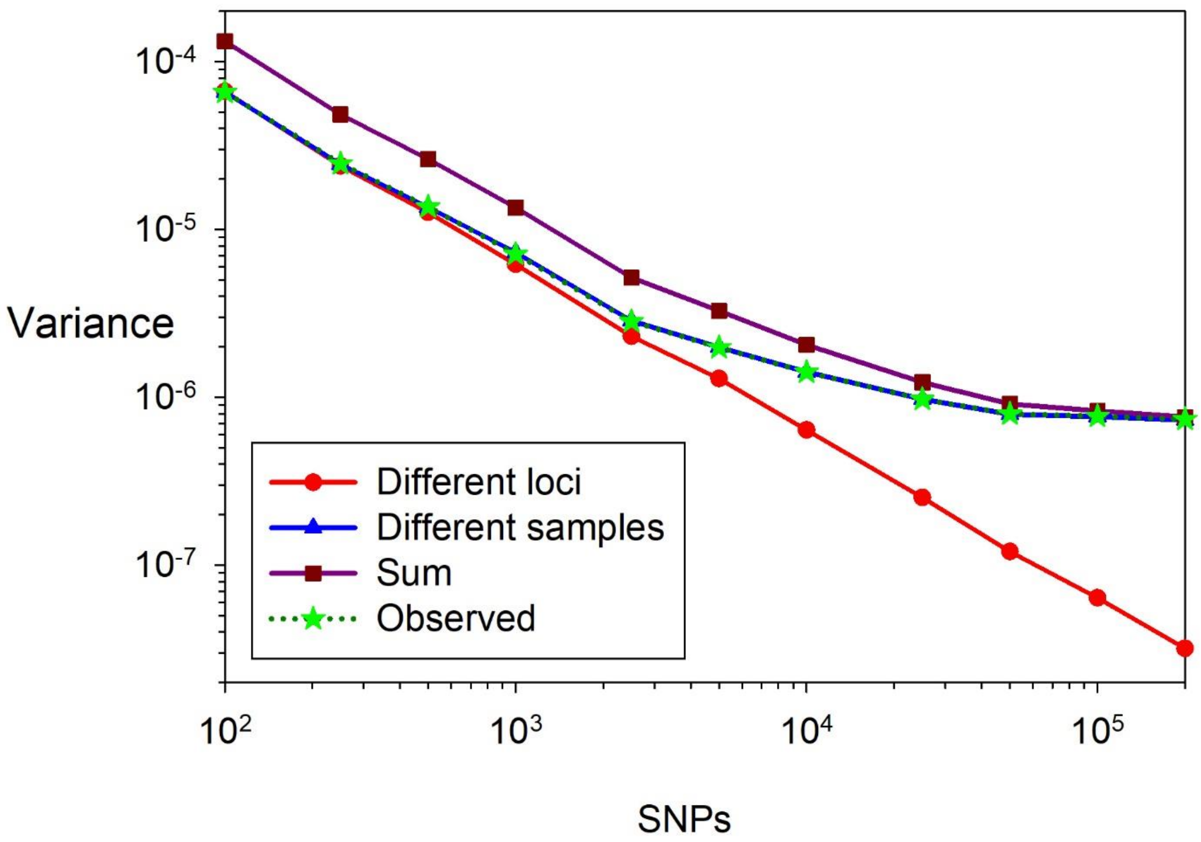


Figure S19. Variance components analysis for $F_{ST}$ (see Figure 4 in main text for analogous results for LD). $V_1$ is the variance of mean $F_{ST}$ for the same individuals assayed for different, non-overlapping sets of loci, and $V_2$ is the variance of mean $F_{ST}$ for different (potentially overlapping) sets of individuals assayed for the same loci. "Sum" = $V_1$+$V_2$ and "Observed" is the total observed variance of mean $F_{ST}$. Because variance associated with sampling individuals dominates, the "Observed" line is largely indistinguishable from the line for $V_2$. Results are for simulations with $N_e$ = 200, $C$ = 16, $S$ = 50.

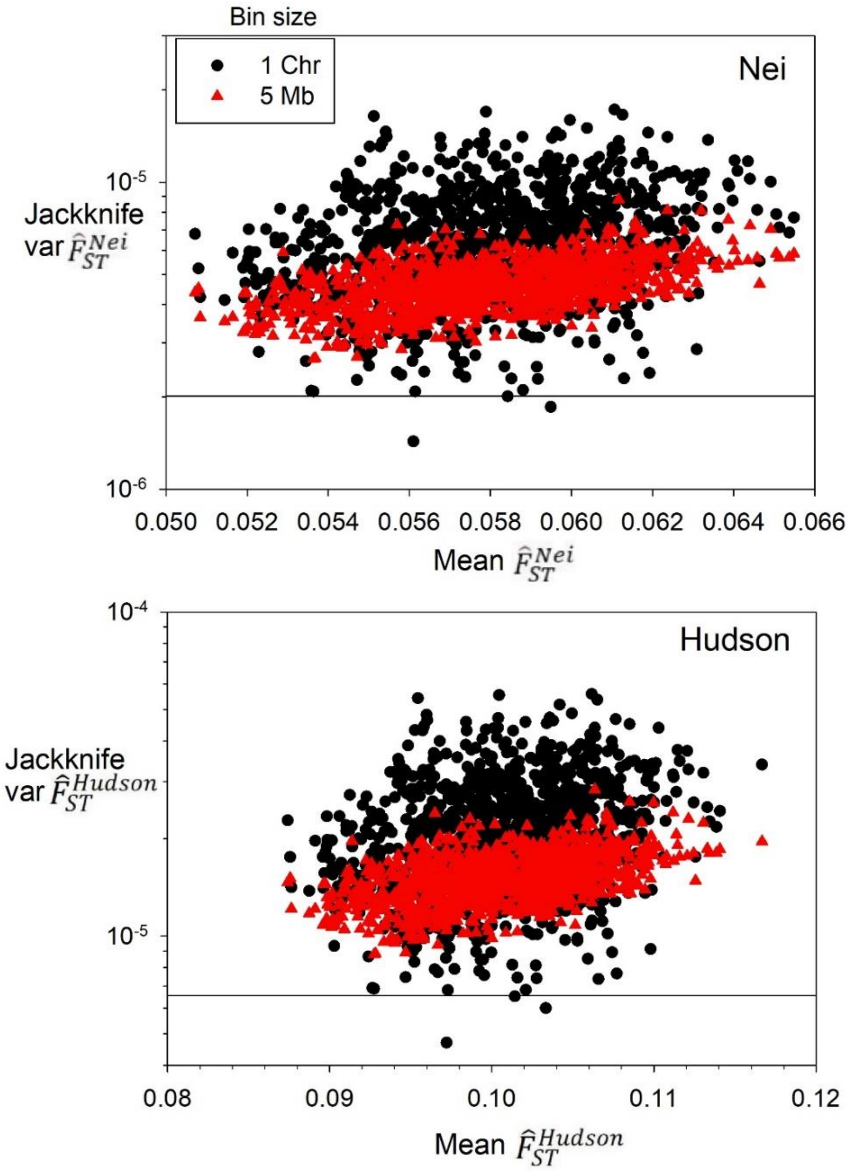


Figure S20. Block jackknife estimates of var($\hat{F}_{ST}^{Nei}$) (top) and var($\hat{F}_{ST}^{Hudson}$) (bottom) for simulations with $N_e$ = 200, $C$ = 16, $S$ = 50, and $L$ = 5000 SNP loci. Black circles show results for single-chromosome blocks and red triangles results for blocks of 5 Mb. Solid horizontal lines show estimates of true var($\hat{F}_{ST}^{Nei}$) and var($\hat{F}_{ST}^{Hudson}$) calculated in this study.
